# Phylogenetic and functional characterization of Asgard primases

**DOI:** 10.1101/2025.10.19.683342

**Authors:** Zhimeng Li, Yang Liu, Li Huang, Meng Li

## Abstract

Eukarya resemble Archaea in DNA replication. Analysis of the DNA replication machinery of Asgard archaea may provide a valuable test of the hypothesis of this phylum being the origin of Eukarya. Among the replication proteins, primase, which comprises the catalytic subunit PriS and the noncatalytic subunit PriL, synthesizes primers for extension by DNA polymerase. Here we show that Asgard primases fall into two major groups, denoted the Heimdall group and the Loki group, which are phylogenetically and structurally more closely related to eukaryotic primases and primases from non-Asgard archaea, respectively. Notably, like human PriL, PriL of the Heimdall group possesses an extra C-terminal domain, which, absent in archaeal PriL of the non-Heimdall group, presumably serves to enhance the stability of the conserved iron-sulfur cluster in PriL. We overproduced in Escherichia coli and purified the PriS and PriL subunits of the Heimdall group from the *Candidatus* Gerdarchaeota archaeon B18_G1. Biochemical characterization reveals that the B18_G1 primase is capable of primer synthesis and extension, using preferentially dNTPs as the substrates, as shown for primases from non-Asgard archaea, but, unlike the non-Asgard archaeal primases, it produces short primers, a feature typical of eukaryotic primases. These results shed significant light on the evolutionary pathway of primase, and are consistent with the hypothesis of the Asgard origin of Eukarya.

## Introduction

The origin of Eukarya has long been of great interest in biology. The endosymbiotic hypothesis posits that the first eukaryotic cell arose from a symbiotic association between an archaeal host cell and an α-proteobacterium, the precursor of mitochondria (Koonin 2015; Staley and Caetano-Anolles 2018; Nobs, et al. 2022). Among the early pieces of evidence in support of the endosymbiotic hypothesis is the striking resemblance of Eukarya to Archaea, and not Bacteria, in DNA replication machinery (Huet, et al. 1983). The discovery of Asgard archaea, which encode numerous eukaryotic signature proteins (ESPs), a decade ago has provided powerful data further strengthening this theory (Koonin 2015; Spang, et al. 2015; Dey, et al. 2016; Surkont and Pereira-Leal 2016; Zaremba-Niedzwiedzka, et al. 2017; Spang, et al. 2018; MacLeod, et al. 2019; Imachi, et al. 2020; Liu, et al. 2021; Imachi, et al. 2025; Zhang, et al. 2025). Five phyla of Asgard archaea, i.e., Lokiarchaeota, Thorarchaeota, Odinarchaeota, Heimdallarchaeota and Helarchaeota, were identified initially (Spang, et al. 2015), and the remarkable expansion of the Asgard family has since ensued (Cai, et al. 2021; Farag, et al. 2021; Liu, et al. 2021). In recent years, some Asgard strains (e.g. ‘*Candidatus* Prometheoarchaeum syntrophicum’ strain MK-D1) have been successively isolated (Imachi, et al. 2020; Imachi, et al. 2025). Asgard archaea are recently reclassified as a single phylum, named Asgardarchaeota or Prometheoarchaeota, within the domain Archaea (Imachi, et al. 2024). Phylogenetic and phylogenomic analyses, as well as phenotypic studies, have led to the speculation that Eukarya may have originated from Asgard archaea (Spang, et al. 2015; Eme, et al. 2023; Zhang, et al. 2025).

Among DNA replication proteins, primases from Archaea and Eukarya are the least closely related (Frick and Richardson 2001; Pellegrini 2012). Primase synthesizes primers for extension by DNA polymerase during DNA replication (Frick and Richardson 2001). Primases are grouped into two major categories, i.e., the single-subunit bacterial primase DnaG and the multi-subunit archaeo-eukaryotic primase (AEP) employed by archaea and eukaryotes (Frick and Richardson 2001; Pellegrini 2012). Eukaryotic primase comprises a heterodimer of the catalytic (small) subunit p49/Pri1/PriS and the non-catalytic (large) subunit p58/Pri2/PriL in complex with DNA polymerase Polα/p180 and the B subunit/p70 (Santocanale, et al. 1993; Copeland 1997; Schneider, et al. 1998; Pellegrini 2012; Suwa, et al. 2015). On the other hand, archaeal primase contains only the catalytic subunit PriS, the non-catalytic subunit PriL and, in some crenarchaea, the second noncatalytic subunit PriX (Desogus, et al. 1999; Liu, et al. 2001; Lao-Sirieix and Bell 2004; Nunez-Ramirez, et al. 2011).

The catalytic core of canonical PriS contains three conserved motifs, i.e., the hhhDhD/E motif (motif I, where “h” represents a hydrophobic residue), the sxH motif (motif II, where “s” stands for small residue, and “x” represents any residue), and the hD/E motif (motif III) (Iyer, et al. 2005). However, archaeal primases exhibit significant sequence divergence across different phyla. The PriS proteins from different archaeal phyla share low amino acid sequence similarity, whereas eukaryotic primases are more evolutionarily conserved (Liu, et al. 2001). As can be expected from their sequence diversity, primases from different archaeal lineages and eukaryotes vary in biochemical activity and physiological function (Frick and Richardson 2001; Pellegrini 2012; O’Donnell and Kurth 2013; Guilliam, et al. 2015). For example, primase from *Saccharolobus solfataricus* P2 exhibits primer synthesis, extension and terminal transfer activities, and synthesizes either DNA or RNA primers *in vitro* (De Falco, et al. 2004; Lao-Sirieix and Bell 2004; Lao-Sirieix, Pellegrini, et al. 2005; Liu, et al. 2015; Yan, et al. 2018). Primase from *Pyrococcus furiosus* preferentially uses dNTPs over rNTPs as the substrates, being able to synthesize DNA products of several kilobases in size (Liu, et al. 2001). By comparison, eukaryotic primases are known to initiate primer synthesis at specific sites, which are not found in archaeal genomes, and synthesize 9-12-nt RNA primers (Frick and Richardson 2001; Baranovskiy, et al. 2016). Therefore, sequence, structural and biochemical analyses of primases from different lineages will shed light on the evolution of primase and provide clues to the origins of both eukaryotic primase and Eukarya.

In this study, we show that Asgard primases fall into two groups, which cluster with typical archaeal primases and eukaryotic primases, respectively. We further demonstrate that primase from *Candidatus* Gerdarchaeota archaeon B18_G1, a representative of primases of the latter group, has unique biochemical properties, which appear to position the enzyme in the evolutionary pathway leading to eukaryotic primase.

## Materials and Methods

### Phylogenetic analysis

Amino acid sequences of priL and priS genes were retrieved from representative archaeal genomes with >70% completeness and <10% contamination in the GTDB database (version R220) (Parks, et al. 2021) using the TADA pipeline (sample_gtdb) (Hägglund, et al. 2023). Target proteins were identified by performing Pfam HMM searches using the domain profiles PF04104 (PriL) and PF01896 (PriS) against the selected proteomes, applying built-in significance cutoffs for domain detection (Mistry, et al. 2020). The resulting alignments were subsequently refined using trimAl (Capella-Gutiérrez, et al. 2009) with the “gappyout” option to remove poorly aligned regions and gap-rich sites that could introduce phylogenetic noise. Phylogenetic reconstruction was performed using IQ-TREE v2 with a two-step approach (Kalyaanamoorthy, et al. 2017; Minh, et al. 2020). Initially, a guide tree was constructed under the LG+F+G substitution model, which accounts for amino acid frequencies and gamma-distributed rate heterogeneity. Subsequently, the second phylogenetic analysis was performed using the LG+C60+G+F+PMSF model, which incorporates a mixture of 60 profile classes with gamma-distributed rate variation, empirical amino acid frequencies, and posterior mean site frequency approximation. The analysis utilized the guide tree from the initial step and employed both bootstrap support (1000 replicates) (Hoang, et al. 2018) and SH-aLRT (Shimodaira-Hasegawa approximate likelihood ratio test, 1000 replicates) (Shimodaira and Hasegawa 1999) to assess branch confidence.

### Structure alignment

Structural alignment and visualization of primases were performed using the Pairwise Structure Alignment tool on RCSB PDB (Sehnal, et al. 2021; Bittrich, et al. 2024). The TM-align method, a sequence-independent structural alignment algorithm (Zhang and Skolnick 2005), was employed for the alignment. Secondary structure alignment was performed with ESPript 3.0 (https://espript.ibcp.fr/ESPript/ESPript/) (Robert and Gouet 2014).

### Preparation of wild-type and mutant primase subunits

DNA fragments containing primase genes from *Candidatus* Gerdarchaeota archaeon B18_G1 (i.e., *Gerd_priS*, *Gerd_epriL* and *Gerd_apriL1*) were synthesized at Beijing Liuhe BGI Co., China, and cloned into plasmid pET30a between the *Nde*I and *Xho*I sites, yielding pET30a-PriS, pET30a-ePriL and pET30a-aPriL1. To coproduce Gerd_PriS and Gerd_ePriL or Gerd_aPriL1, the two corresponding encoding genes were inserted between *Nde*I and *Kpn*I, and between *Not*I and *Xho*I, respectively. Mutant Gerd_PriS proteins (i.e., Y86F and Y86D) were obtained from pET30a-PriS by using the Fast Mutagenesis System kit (Transgen Biotech, Beijing, China). *E. coli* BL21(AI) cells was transformed with the plasmids and cultured at 37°C with shaking at 200 rpm to an OD_600_ of ∼0.5 in LB medium containing 50 μg/ml kanamycin. The culture was then incubated at 16°C for 1 hr. Induction of the protein synthesis was achieved by the addition of L-arabinose and IPTG to final concentrations of 0.2% and 0.2 mM, respectively. Incubated continued for overnight at 16°C with shaking. The cells were harvested and frozen at −80°C.

To purify recombinant proteins, the cells were lysed by sonication, and the lysate was incubated at 55°C (45°C for the lysate of the Y86D overproducer) for 10 min and centrifuged at 17,000 x g to remove bulk of the *E. coli* proteins. The clarified lysate was treated 0.5% polyethylenimine (pH 7.9) and centrifuged. Ammonium sulfate was added to the supernatant to 436 mg/ml. After centrifugation, the precipitate was collected, resuspended in buffer A [50 mM Tris-HCl, pH 6.8, 150 mM NaCl, 15 mM imidazole, 10% (v/v) glycerol], and loaded onto a Histrap HP column (Cytiva, USA), equilibrated with buffer A, on an AKTA Pure system (Cytiva, USA). Proteins were eluted with a 0-100% gradient of buffer B (i.e., buffer A containing 500 mM imidazole). Fractions containing the target protein, which came off the column at 50-75 mM imidazole, were collected, pooled and dialyzed against buffer C [50 mM Tris-HCl, pH 6.8, 50 mM NaCl, and 10% (v/v) glycerol]. The dialysate was loaded onto a HiTrap Heparin HP column (Cytiva, USA), equilibrated with buffer C, on the AKTA system. Elution was carried out with a 0-100% gradient of buffer D [50 mM Tris-HCl, pH 6.8, 1 M NaCl, and 10% (v/v) glycerol]. Proteins, eluted at 150-250 mM NaCl, were concentrated by ultrafiltration (Millipore, USA), and further purified on a Superdex 200 column (Cytiva, USA) in buffer E [50 mM MES-NaOH, pH 7.0, 150 mM NaCl, and 15% (v/v) glycerol]. Fractions containing the target protein were collected and pooled. The concentrations of the purified proteins were determined by the Lowry method using bovine serum albumin (BSA) as the standard. The proteins were stored at −80°C.

Recombinant primases from *Pyrococcus furiosus* and *Sulfolobus solfataricus* P2 (Table S1) were prepared and purified as described previously (Liu, et al. 2001; Liu, et al. 2015).

### Surface plasmon resonance (SPR)

The SPR assays were performed on a BIACORE 3K system (Cytiva, USA). Purified Gerd_ePriL or Gerd_aPriL1 was immobilized on CM5 sensorchips (Cytiva, USA). All experiments were conducted in buffer containing 20 mM Hepes, pH 7.5, 500 mM NaCl, and 0.005% Tween 20 and flowing at a rate of 30 μl/min at 25°C. The dissociation constants of the Gerd_PriS/ePriL and Gerd_PriS/aPriL1 complexes were determined with a Gerd_PriS concentration gradient.

### Primer synthesis assays

The standard reaction mixture (10 μl) contained 1.5 μM primase, 230 ng M13mp18 ssDNA or 1 μM dT_35_ or SP2 (Table S1), 10 μM dNTPs (1 μCi [α-^32^P]dATP) and 100 μM rNTPs in 50 mM MES-NaOH (pH 7.0), 100 μg/ml BSA, and 10 mM MnCl_2_. After incubation for 30 min at 55°C, the reactions were stopped and deproteinized as described in the primer extension assays above. The samples were loaded onto an 25% polyacrylamide gel (19:1) containing 6 M urea. After electrophoresis in 1 x TBE buffer, the gel was exposed to X-ray film.

### Primer extension assays

The standard reaction mixture (10 μl) contained 50 mM MES-NaOH, pH7.0, 100 μg/ml BSA, 10 mM MnCl_2_, 10 μM rNTPs/dNTPs, 1.5 μM primase, and 4 nM labeled template. The labeled template was prepared by annealing a labeled primer (D25 or R25, Table S1) with M13mp18 ssDNA (New England Biolabs, USA) at 1:1.25 molar ratio. D25 and R25 were labeled at the 5’ terminus using [γ-^32^P]ATP and T4 polynucleotide kinase (New England Biolabs, USA). The primer extension reaction was incubated for 30 min at 55°C. The SDS and proteinase K were added to the final concentrations of 0.8% and 1.6 mg/ml, respectively. After incubation at 55°C for 3 hrs, the mixture was extracted with an equal volume of phenol-chloroform-isoamyl alcohol (25:24:1) and the nucleic acid precipitated with ethanol. The precipitate was resuspended in loading buffer (98% formamide, 10 mM EDTA, 0.025% xylene blue and 0.025% bromophenol blue), boiled for 5 min, and cooled rapidly on ice. The sample was subjected to electrophoresis in 15% polyacrylamide gel (19:1) containing 8 M urea in 1 x Tris-borate-EDTA (TBE) buffer. The gel was exposed to X-ray film.

### Electrophoretic mobility shift assays (EMSA)

A 25-nt single-stranded DNA fragment (D25) of the same sequence as the primer used in the extension assays was labeled at the 5’ terminus using [γ-^32^P]ATP and T4 polynucleotide kinase (New England Biolabs, USA). Primase (25 nM-2 μM) was incubated for 15 min at 23°C with 4 nM DNA template in 50 mM MES-NaOH, pH 7.0, 100 μg/ml BSA, 0.05% Triton X100, and 8% (v/v) glycerol in a total volume of 10 μl. The samples were loaded on an 8% native polyacrylamide gel. Following electrophoresis in 0.2 x TBE buffer, the gel was exposed to X-ray film.

### Thermal stability

Aliquots (10 μl) of the primase preparation (10 μM) were incubated for 15 min at indicated temperatures, and centrifuged at 17,000 x g for 10 min to remove the precipitate. After the addition of the loading buffer (10 μl) for SDS-PAGE, the mixtures were heated for 5 min in boiling water and cooled rapidly in ice. Following centrifugation at 17,000 x g for 5 min, the supernatants were subjected to 12% SDS-PAGE.

## Results

### Phylogenetic analysis of primases

To elucidate the evolutionary relationship of archaeal-eukaryotic primases, we constructed comprehensive phylogenetic trees for the primase subunits PriS and PriL. The analysis incorporated 232 PriS sequences, including representatives from major archaeal lineages (89 Asgard, 8 DPANN, 37 Thermoproteota, 37 Thermoplasmatota, 17 Halobacteriota, 15 Methanobacteriota, 4 Hadarchaeota, and 3 Hydrothermarchaeota sequences) and 22 eukaryotic sequences. For PriL, we analyzed 226 sequences with comparable taxonomic representation (77 Asgard, 9 DPANN, 39 Thermoproteota, 36 Thermoplasmatota, 15 Halobacteriota, 15 Methanobacteriota, 4 Hadarchaeota, 3 Hydrothermarchaeota, and 28 eukaryotic sequences), ensuring coverage across all known archaeal phyla at the class or family level.

The observed phylogenetic patterns reveal that primases from Asgard archaea segregate into two evolutionarily distinct groups showing different relationships with eukaryotic primases (Fig. 1A, 1B, S1A, and S1B). In the PriL tree, eukaryotic sequences formed a sister group to a subset of Asgard lineages containing predominantly Heimdallarchaeia, including Hodarchaeales, Heimdallarchaeales, and Gerdarchaeales, along with Wukongarchaeia, Njordarchaeia, and Sifarchaeia (Borrarchaeales). This group is designated as the Heimdall group or Asgard-eukaryotic primase clade. Other Asgard lineages, including Odinarchaeia, Lokiarchaeia, Jordarchaeia, Thorarchaeia, and Hermodarchaeia, cluster with Thermoproteota rather than eukaryotes. This group is named the Loki group or Asgard-archaeal primase clade. The PriS phylogeny shows eukaryotic sequences nested within the Asgard clade comprising Heimdallarchaeia, Wukongarchaeia, Njordarchaeia and Sifarchaeia, with the remaining Asgard lineages again clustering with Thermoproteota. In addition, Panguiarchaeum, which was previously considered to belong to the TACK superphylum (Qu, et al. 2023) and recently shown to be an Asgard archaeon (Huang, et al. 2025), also appears in the Heimdall group.

**Figure 1.**
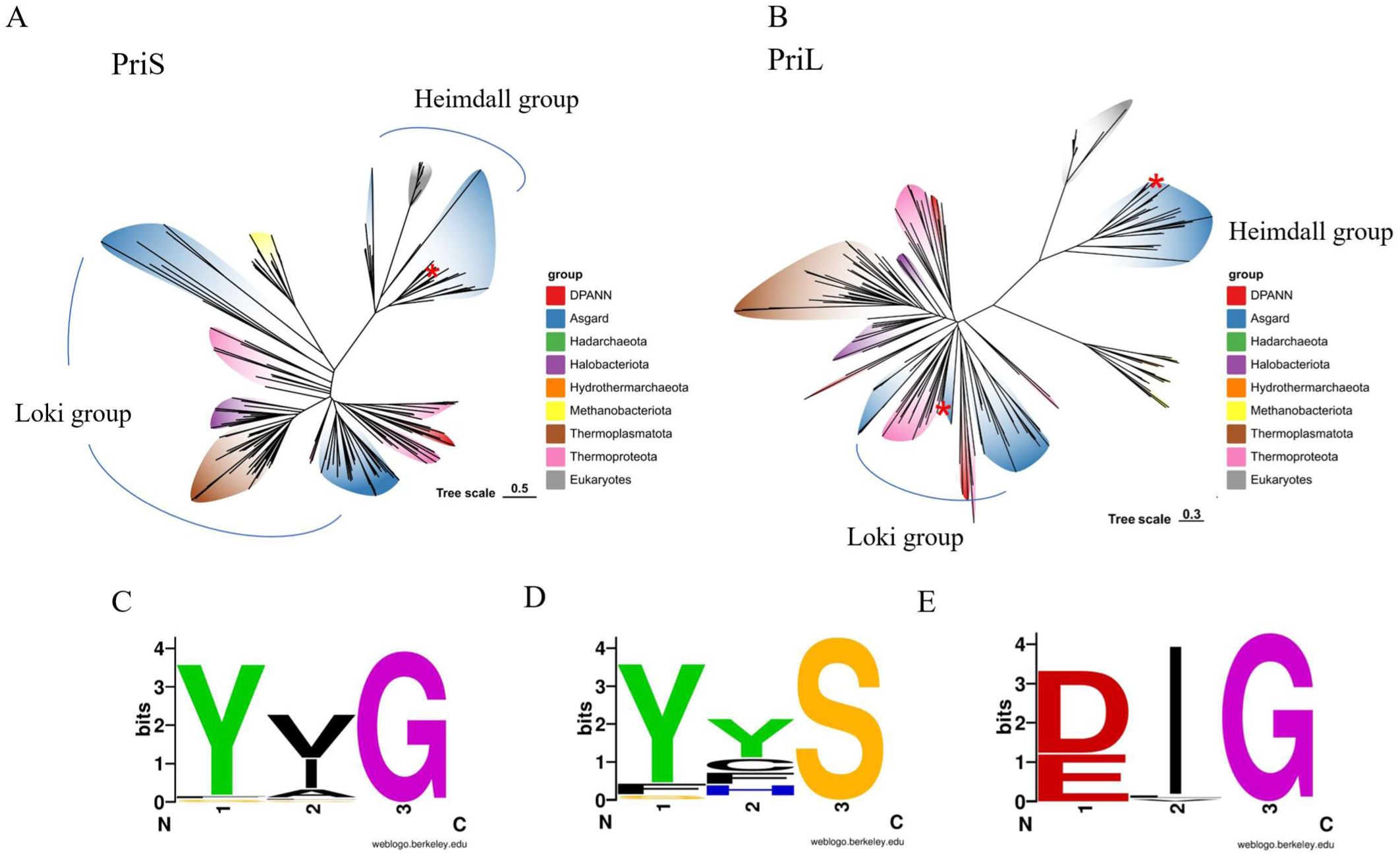
Phylogenetic analysis of primases from Archaea and Eukarya. **(A and B)** Maximum likelihood phylogeny trees of archaeal and eukaryotic PriS (A) and PriL (B). The Heimdall and Loki groups of Asgard primases are marked with arcs. *, Location of primases encoded by *Candidatus* Gerdarchaeota archaeon B18_G1 **(C, D and E)**. The sequence logo of motif G/S in PriS from Asgard primases of the Heimdall group (C), remaining Archaea (D) and Eukarya (E). The height of a letter represents the strength of conservation. The logo was generated via weblogo.berkeley.edu (https://weblogo.berkeley.edu/logo.cgi).

Sequence analysis of the PriS subunits from the Heimdall group and eukaryotes identifies a conserved glycine (G) residue approximately 25 amino acids (aa) upstream of the hhhDhD/E motif (motif I). This position, typically occupied by a serine (S) residue in PriS from non-Heimdall archaea, along with the two preceding amino acid residues will be referred to herewith as motif G/S. The sequence of motif G/S is generally Y*G (Fig 1C) (“*” represents a variable residue), with tyrosine occasionally replaced by phenylalanine (F), in primases of the Heimdall group, and Y/F/S*S, with “*” being highly variable, in other archaeal groups (Fig 1D). This motif in eukaryotic primases is often D/E*G, with E being almost exclusively present in fungal PriS (Fig 1E).

### Primase from *Candidatus* Gerdarchaeota archaeon B18_G1

To gain insight into the biochemical properties of Asgard primase, we attempted to overproduce primase subunits from Lokiarchaeia, Thorarchaeia, Gerdarchaeales and Wukongarchaeia, which represent primases from both the Heimdall and Loki groups as well as with different types of motif G/S. However, despite considerable efforts, we only succeeded in overexpressing primase genes from the metagenome-assembled genome (MAG) of *Candidatus* Gerdarchaeota archaeon B18_G1, which was derived from a deep-sea hydrothermal vent sediment, in *E. coli* (Fig. S2A). The B18_G1 genome encodes one PriS subunit (B18_G1_heimdall_00653) and three PriL subunits (B18_G1_heimdall_00652/02569/02778). B18_G1_heimdall_00653 belongs to the Heimdall group (denoted Gerd_ePriS) with motif G/S being YVG. Intriguingly, of the three PriL subunits, B18_G1_heimdall_00652 is in the Heimdall group (Gerd_ePriL), while B18_G1_heimdall_02569 and B18_G1_heimdall_02778 belong to the Loki group (Gerd_aPriL1 and Gerd_aPriL2, respectively). Recombinant Gerd_ePriS, Gerd_ePriL and Gerd_aPriL1 were purified, but recombinant Gerd_aPriL2 turned out to be insoluble.

Genes encoding PriS and PriL are adjacently situated in the genomes of all Heimdall lineages except for Borrarchaeales and some Hodarchaeales (Fig. S1). In the genome of B18_G1, the Gerd_ePriL gene, but not the genes for Gerd_aPriL1 and Gerd_aPriL2, resides next to the Gerd_ePriS gene. Based on this observation as well as clustering of the Gerd_aPriL1 and Gerd_aPriL2 genes with PriL sequences from Bathyarchaeia in the phylogenetic trees, we speculate that the Gerd_ePriL gene is vertically inherited, while the genes encoding Gerd_aPriL1 and Gerd_aPriL2 may have acquired through horizontal gene transfer.

### Structure comparison of primases

We then predicted the 3-D structures of B18_G1 primase subunits using AlphaFold and compared them with the experimentally resolved or AlphaFold-predicted structures of human (Baranovskiy, et al. 2015), *Saccharolobus solfataricus* (Lao-Sirieix, Nookala, et al. 2005; Holzer, et al. 2017), *Promethearchaeum syntrophicum* (Imachi, et al. 2020; Imachi, et al. 2024) and *Pyrococcus furiosus* (Augustin, et al. 2001) primases (Fig. S3, Table S2 and S3). Among these primases, those from *P. furiosus* and *S. solfataricus* are more closely related to the Heimdall and the Loki groups, respectively. All PriS subunits share a conserved α/β Prim fold, in which β-sheets form a groove where the catalytic center is located (Fig. 2A, S3A and S4) (Augustin, et al. 2001; Lao-Sirieix, Nookala, et al. 2005; Kilkenny, et al. 2013; Baranovskiy, et al. 2015). Structural differences among these PriS subunits occur primarily in the peripheral regions of the proteins (Fig. 2A and S3A). Pairwise structural comparison shows that Gerd_ePriS is most similar to human PriS (TM-score: 0.81), followed by *P. furiosus* PriS (0.74), *P. syntrophicum* PriS (0.62) and *S. solfataricus* PriS (0.62) (Table S4). It is worth noting that distinctive differences were observed between Gerd_ePriS and *P. syntrophicum* PriS, which belong to the Heimdall and Loki groups, respectively, although both of them are from Asgard archaea (Fig. S4).

**Figure 2.**
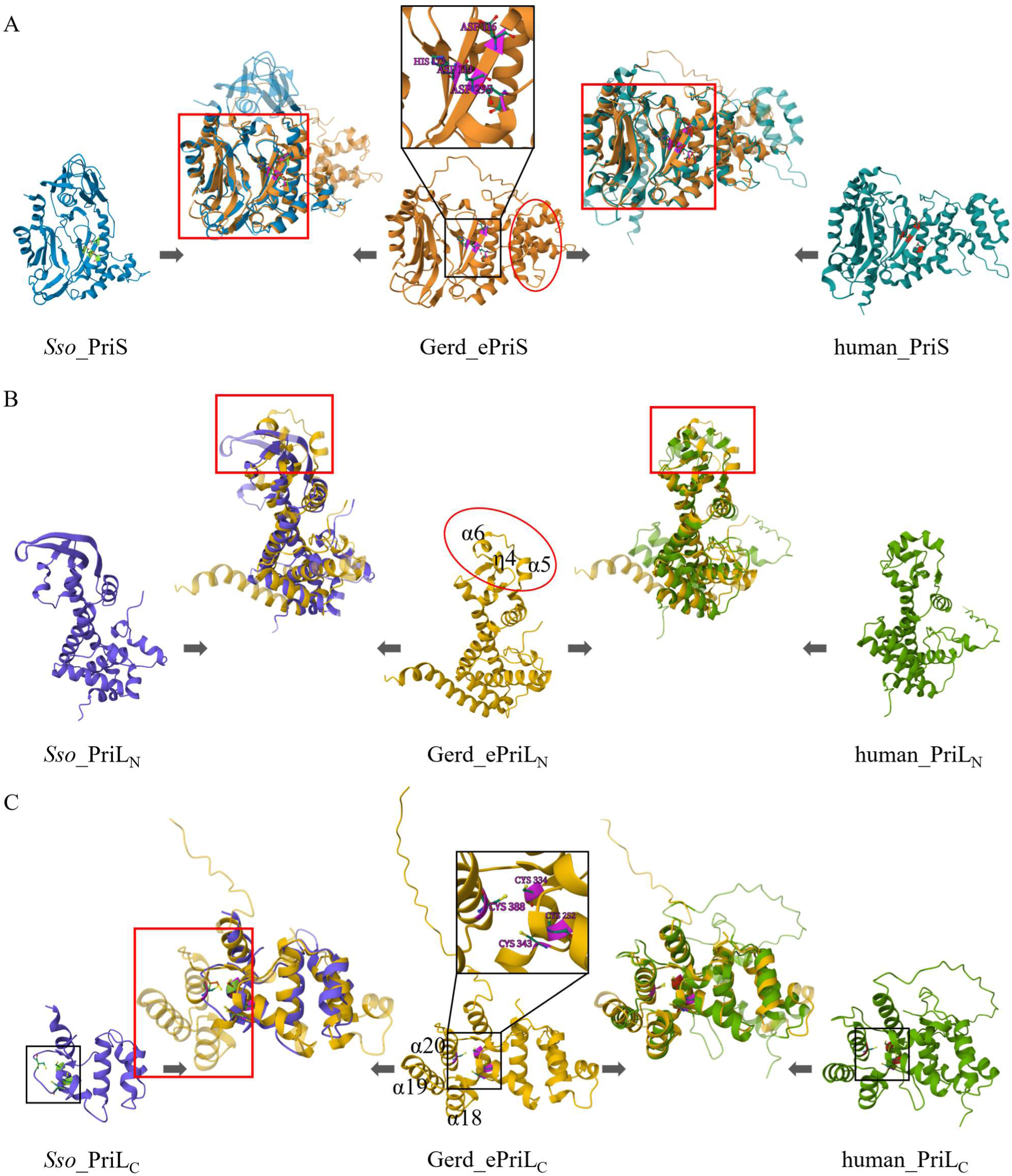
Structural alignment of primases. AlphaFold-predicted models of *Candidatus* Gerdarchaeota archaeon B18_G1 and *Promethearchaeum syntrophicum* primases were downloaded from the AlphaFold Database (https://alphafold.com/). Experimentally resolved structures of human (PDB: 4MHQ & 5EXR), *Saccharolobus solfataricus* (PDB: 5OF3), and *Pyrococcus furiosus* (PDB: 1g71) were retrieved from RCSB PDB (https://www.rcsb.org/). A portion of PriL-CTD has yet to be resolved for any PriL proteins and was predicted with AlphaFold (Table S2). The structure of *P. furiosus* PriL was derived from the partial crystal structures of its homologues from *Pyrococcus abyssi* and *Pyrococcus horikoshii* via AlphaFold (Table S2). **(A)** PriS. The Prim fold domain and the catalytic core of Gerd_ePriS are shown in red and black boxes, respectively. The key amino acid residues (D114, D116, H173 and D295) at the catalytic center are indicated. The region comprising β9 to α11 at the PriSL interface is marked with a red oval. **(B)** PriL-NTD. The region comprising α5, α6 and η4 at the PriSL interface is marked with a red oval. The region used in the structural alignment is shown in red box. **(C)** PriL-CTD. The location of the iron-sulfur clusters is shown in black box. The four cysteine residues (C252, C334, C343 and C388) of the iron-sulfur cluster in Gerd_ePriL are indicated. The C-terminal domain with three α-helices (α18, α19 and α20 in Gerd_ePriL), which is unique to eukaryotic and the Heimdall group PriL proteins, is shown in black box. *Sso*, *S. solfataricus*; Gerd, *Candidatus* Gerdarchaeota archaeon B18_G1; PriL_N_, the N-terminal domain of PriL; PriL_C_, the C-terminal domain of PriL. The prefix ‘e’ in ePriL_N_ and ePriL_C_ denotes the protein of the Heimdall group. Regions of identity and difference between two aligning proteins are shown as semitransparent and opaque, respectively. Structural alignments were performed using the Pairwise Structure Alignment tool on RCSB PDB with TM-align method.

Since PriL proteins are not amenable to direct structural alignment due to the presence of two structural domains (N- and C-terminal domains, or NTD and CTD) connected with a highly flexible linker region, we compared these proteins at the domain level. The overall structures of NTDs from different PriL proteins are highly similar, with the most divergent region corresponding to residues ∼120-140 in Gerd_ePriL (α5, α6, and η4) (Fig. 2B, S3B and S5). Adjacent to this region in the PriSL complex is a highly divergent region in PriS, which spans from β9 to α11 in Gerd_ePriS (Fig. 2A and S4), raising the possibility that the divergence in the two adjacent regions in the PriSL complex may have resulted from their co-evolution constrained by protein-protein interaction. The CTDs of PriL from different taxa also share a similar architecture. Notably, however, Gerd_ePriL, like human PriL, contains an additional C-terminal domain composed of three α-helices (α18 - 20), which are not present in PriL subunits of the non-Heimdall group (Fig. 2C, S3C and S5). This domain allows the highly conserved iron-sulfur cluster to be enclosed, presumably enhancing its stability. Consistent with this structural feature, our Gerd_ePriL preparation remained brownish, the characteristic color of an iron-sulfur protein containing Fe^2+^, while Gerd_aPriL1, which lacked the C-terminal three α-helices, turned colorless after storage for two weeks at 4°C (Fig. S2B). Gerd_ePriL exhibits the highest similarity to human PriL, among the tested PriLs, in both NTD and CTD with TM-scores of 0.70 and 0.77, respectively (Table S5 and S6). Therefore, our structural alignment results agree with the phylogenetic analysis in suggesting that Asgard primases fall into two major groups, with the Heimdall group being more evolutionarily closely related to eukaryotic primases than other archaeal primases.

### Gerd_ePriS binds to Gerd_ePriL more tightly than to Gerd_aPriL

Since B18_G1 encodes a single Heimdall group PriS protein (Gerd_ePriS) but PriL proteins of both the Heimdall and Loki types (Gerd_ePriL and Gerd_aPriL1), we first sought to determine which PriL might form a complex with the PriS by SPR. As shown in Figs. 3A and 3B, Gerd_ePriS bound far more tightly to Gerd_ePriL (dissociation constant [KD]: 2.667 ± 1.204 x 10^−9^ M) than to Gerd_aPriL1 (KD: 1.054 ± 0.196 x 10^−7^ M). The stability of the Gerd_ePriS/ePriL complex was evidenced by the failure to elute the bound Gerd_ePriS off the immobilized Gerd_ePriL with buffer containing 2 M NaCl (data not shown). By comparison, bound Gerd_ePriS was readily washed off the immobilized Gerd_aPriL1 at the same salt level. We then conducted a competition experiment in which a mixture of Gerd_ePriS and Gerd_ePriL and that of Gerd_ePriS and Gerd_aPriL1 were separately flown through the chip immobilized with Gerd_aPriL1. In a control experiment, when flown through alone, Gerd_ePriS was able to bind to Gerd_aPriL1 on the chip. A binding curve similar to that with Gerd_ePriS alone was obtained when Gerd_ePriS-aPriL1 was flown through, suggesting that immobilized Gerd_aPriL1 was able to bind or grab Gerd_ePriS from the complex (Fig. 3D). In comparison, binding of Gerd_ePriS to immobilized Gerd_aPriL1 did not occur when Gerd_ePriS-ePriL was flown through (Fig. 3C). These results suggest that Gerd_ePriS forms primarily a complex with Gerd_ePriL, and not with Gerd_aPriL1, in the cell.

**Figure 3.**
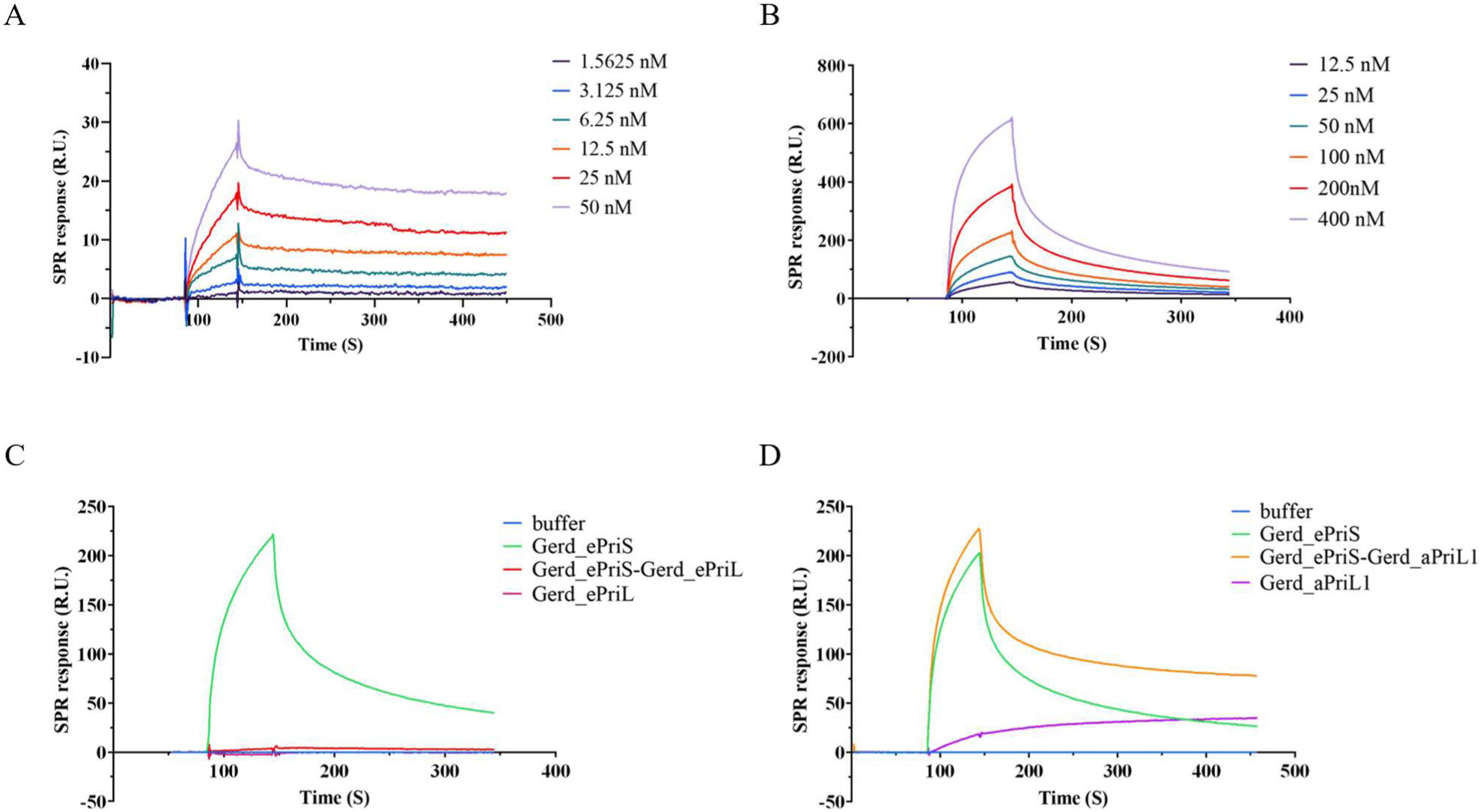
Analysis of the interactions of Gerd_ePriS with Gerd_ePriL and Gerd_aPriL1 by SPR. **(A and B)** Interaction of Gerd_ePriS with Gerd_ePriL and Gerd_aPriL1. Gerd_ePriL (A) or Gerd_aPriL1 (B) flowed through the chip immobilized with Gerd_ePriS as the mobile phase. Gradients of protein concentrations used in the experiments are indicated. **(C & D)** Binding competition assays. A mixture of Gerd_ePriS and Gerd_ePriL **(C)** or that of Gerd_ePriS and Gerd_aPriL1 **(D)** flowed through the chip immobilized with Gerd_aPriL1 as the mobile phase. Elution buffer, Gerd_ePriL or Gerd_aPriL1 are the negative controls, and Gerd_ePriS is the positive control.

### Gerd_ePriSL synthesizes short primers

We then examined the ability of B18_G1 primase (Gerd_ePriSL) to synthesize primers. Gerd_ePriS exhibited very low but detectable primer synthesis activity, when rNTPs were used as the substrates, even in complex with Gerd_ePriL or Gerd_aPriL1, on the M13mp18 ssDNA template (Fig. S6). Slightly more primer synthesis by Gerd_ePriS or its complex with Gerd_ePriL or Gerd_aPriL1 was observed in the presence of dNTPs instead of rNTPs (Fig. S6). Notably, primer synthesis by Gerd_ePriSL increased significantly in the presence of both dNTPs and rNTPs at the dNTPs:rNTPs ratio of 1:1 or, more drastically, 1:10 (Fig. S7 and 4C), which is close to the intracellular rNTP:dNTP ratio (Liew, et al. 2016). Optimal primer synthesis by Gerd_ePriSL occurred at 50∼60^ο^C (Fig. 4B). Among the tested divalent cations, Mn²⁺ was preferred for the activity of the protein (Fig. S8), as found for other archaeal primases (Bocquier, et al. 2001). Notably, primers synthesized by Gerd_ePriSL on M13mp18 ssDNA were very short with sizes mostly no longer than 5 nt (Fig 4A). Slightly longer products (∼9 nt) were synthesized by the enzyme on oligonucleotide templates (Fig. 4A). For comparison, we also determined primer synthesis by primases from *Pyrococcus furiosus* (*Pfu*P41-P46) and *Saccharolobus solfataricus* P2 (*Sso*PriSLX). Both primases synthesized far longer products than Gerd_ePriSL in the presence of either dNTPs or rNTPs or both (Fig. 4C). As shown previously, *Pfu*P41-P46 preferred dNTPs, whereas *Sso*PriSLX was more active with rNTPs in primer synthesis (27,28,30,48). It appears that Gerd_ePriSL is more similar to eukaryotic primases than to the other characterized archaeal primases in terms of the size of their products.

**Figure 4.**
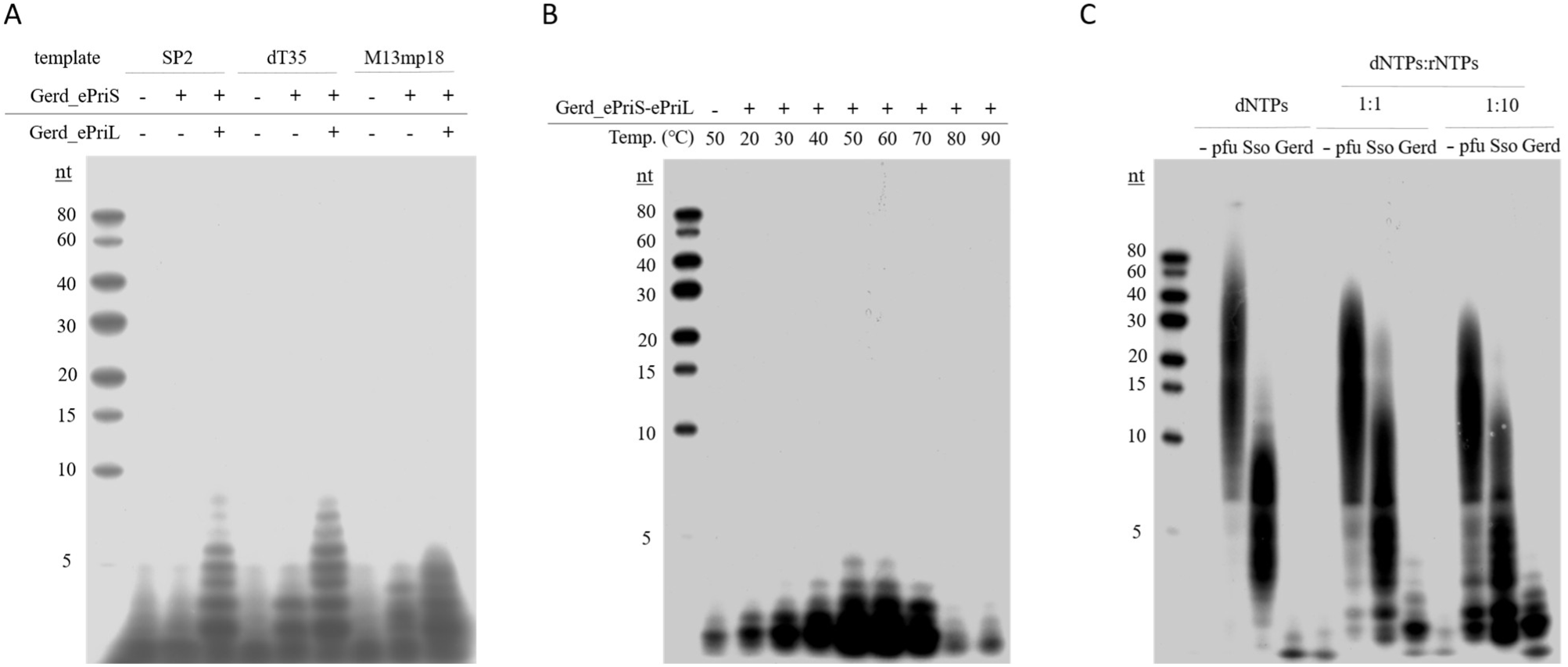
Primer synthesis by B18_G1 primase. **(A)** Primer synthesis on various templates. Protein was incubated with indicated templates (1 μM for SP2 or dT35 and 230 ng for M13mp18 ssDNA), 10 μM dNTPs (1 μCi [α-^32^P]dATP) and 100 μM rNTPs in the standard assay mixture for 30 min at 55℃. **(B)** Primer synthesis at indicated temperatures. Protein (1.5 μM) was incubated with M13mp18 ssDNA (230 ng), 10 μM dNTPs (1 μCi [α-^32^P]dATP) and 100 μM rNTPs in the standard assay mixture for 30 min at indicated temperatures. **(C)** Comparison of primases from *Pyrococcus furiosus* (pfu), *Sulfolobus solfataricus* P2 (*Sso*) and Gerdarchaeota B18_G1 (Gerd) in priming synthesis on different substrates. The above reactions were performed by incubating different primase (1.5 μM) with 230 ng M13mp18 ssDNA and different types of substrates in the standard assay mixture for 30 min at 55℃ for Gerd_ePriSL and *Sso*_PriSLX) or 75℃ for *pfu*_PriSL. dNTPs were added to 10 μM (1 μCi [α-^32^P]dATP), and rNTPs were added to achieve indicated molar ratios. All reactions were stopped by the addition of SDS (0.8%) and protease K (1.6 mg/ml). The products were extracted with phenol/chloroform/isoamyl alcohol (25:24:1) and precipitated with ethanol, and analyzed by 25% polyacrylamide gel (19:1) containing 6 M urea. The gel was exposed to X-ray film.

As mentioned above, motif G/S represents a distinguishing feature of the primase catalytic subunits of various origins. The sequence of this motif in Gerd_ePriS is YVG, FYS in *S. solfataricus* P2 PriS and D*G (*denotes any amino acid) in eukaryotic PriS. The first amino acid residue in motif G/S, located in the groove in close proximity to the catalytic center resides (Fig. S9), was previously suggested to be related to the substrate preference of primase (Li, et al. 2022). In this study, we introduced a point mutation into Gerd_PriS at this position, yielding the mutant proteins Y86F and Y86D, which mimic the first motif G/S residue in PriS from *S. solfataricus* P2 and eukaryotes, respectively. We then measured the thermal stability of the mutant proteins by determining protein precipitation following heat treatment (Fig. 5A). Y86F remained at least as stable as the wild-type protein and was measurably denatured by treatment for 15 min at ≥ 60°C. Interestingly, the thermal stability of eukaryotic-like Y86D was reduced by ∼10°C as compared to the wild-type enzyme, consistent with the proposal that the evolution from Archaea to Eukarya was accompanied by a gradual decrease in optimal growth temperature (Lu, et al. 2024).

**Figure 5.**
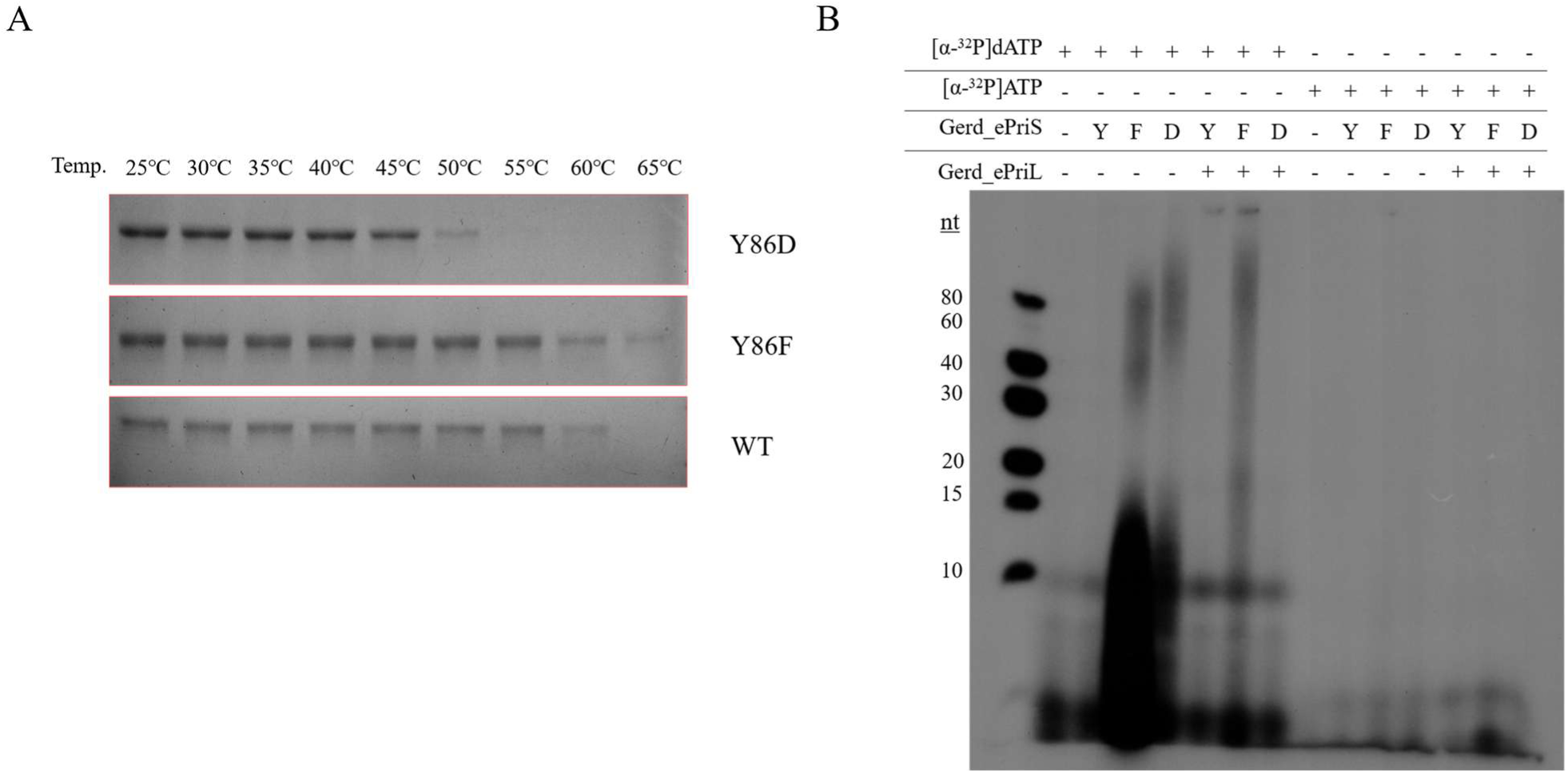
Effect of mutation in motif G/S on the thermostability and primase activity of Gerdarchaeales primase. **(A)** Thermostability of the wild-type and mutant Gerd_ePriS. Proteins (∼ 3 μg) were incubated for 15 min at indicated temperatures, and centrifuged at 17,000 x g for 10 min. The supernatants were subjected to 12% SDS-PAGE. Y86D, Gerd_ePriS_Y86D_; Y86F, Gerd_ePriS_Y86F_; WT, wild-type Gerd_ePriS. **(B)** Primer synthesis by wild-type and mutant Gerd_PriS in the presence and absence of Gerd_ePriL. Reactions were performed by incubating the enzymes (1.5 μM) with M13mp18 ssDNA (230 ng), 10 μM dNTPs and 100 μM rNTPs, 1 μCi [α-^32^P]dATP or 1 μCi [α-^32^P]ATP), as indicated, in the standard assay mixture for 30 min at 45℃ for Gerd_ePriS_Y86D_ or 55℃ for wild-type and Gerd_ePriS_Y86F_. Reactions were stopped by the addition of SDS (0.8%) and protease K (1.6 mg/ml). The products were extracted with phenol/chloroform/isoamyl alcohol (25:24:1), precipitated with ethanol, and analyzed on 25% polyacrylamide gel (19:1) containing 6 M urea. The gel was exposed to X-ray film. Y, Gerd_ePriS; F, Gerd_ePriS_Y86F_; D, Gerd_ePriS_Y86D_.

We then measured the primer synthesis activity of the mutant proteins in the presence of dNTPs:rNTPs at the 1:10 ratio with either [α-^32^P]ATP or [α-^32^P]dATP as the radiolabel at temperatures set according to the thermal stability of the proteins (45°C for Y86D and 55°C for Y86F). Surprisingly, products synthesized by the two mutant proteins far exceeded the wild-type protein in both amount and size (Fig. 5B). The increase in the activity of Y86F was drastically reduced, and that of Y86D was eliminated, by the addition of Gerd_ePriL. Further, both mutant proteins still exhibited the same substrate preference as the wild type protein. Therefore, the two subunits of primase appear to have coevolved to meet the need of the organism.

### Gerd_ePriSL is capable of efficient DNA polymerization

In Eukarya, the PriSL heterodimer synthesizes RNA primers, which are extended by DNA polymerase Polα to generate RNA/DNA hybrid primers (Frick and Richardson 2001; Kilkenny, et al. 2013; Vaithiyalingam, et al. 2014; Baranovskiy, et al. 2015; Guilliam, et al. 2015). In Archaea, by comparison, PriSL possesses polymerase activity and can synthesize long DNA strands (Bocquier, et al. 2001; Liu, et al. 2001; De Falco, et al. 2004; Liu, et al. 2015; Schneider, et al. 2023). To determine if B18_G1 primase carries polymerase activity, we performed primer extension assays using a radiolabeled 25-nt DNA primer annealed to M13mp18 ssDNA as the template. As shown in Fig. 6, Gerd_ePriS alone was capable of extending primer with either dNTPs or rNTPs, and preferentially with the former. The polymerase activity of Gerd_ePriS was optimal at pH 6.5∼9 and 50∼55°C (Fig. S10). This appears to agree with the conditions of the sampling site (deep-sea hydrothermal vent sediments) of the genome B18_G1 (Gerdarchaeales).

**Figure 6.**
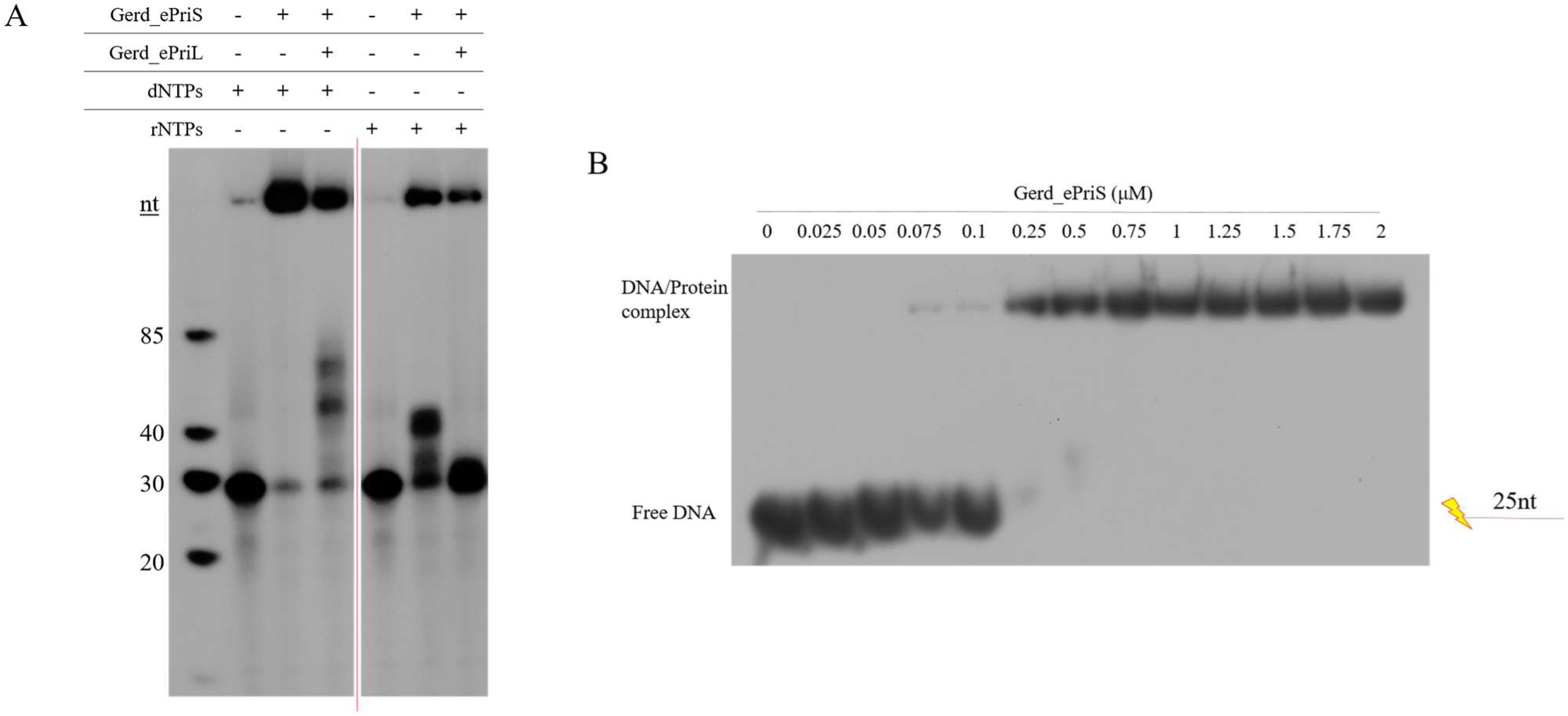
Primer extension by B18_G1 primase and DNA binding by Gerd_ePriS. **(A)** Primer extension by B18_G1 primase. The reaction mixture containing Gerd_ePriS or Gerd_ePriSL (1.5 μM), 4 nM ^32^P-labeled D25 annealed to M13mp18 ssDNA, 10 μM dNTPs (1 μCi [α-^32^P]dATP) or 10 μM rNTPs (1 μCi [α-^32^P]ATP), 50 mM MES-NaOH, pH7.0, 100 μg/ml BSA, and 10 mM MnCl_2_ was incubated at 55℃ for 30 min. Reactions were stopped by the addition of SDS (0.8%) and protease K (1.6 mg/ml). The products were extracted with phenol/chloroform/isoamyl alcohol (25:24:1), precipitated with ethanol, and analyzed on 15% polyacrylamide gel (19:1) containing 8 M urea. The gel was exposed to X-ray film. **(B)** DNA binding by Gerd_ePriS. Gerd_ePriS at indicated concentrations were mixed with 4 nM ssDNA (D25, Table S1), which was labeled with [γ-^32^P]ATP at the 5’-terminus. After 15 min at room temperature, reaction products were analyzed by 8% PAGE followed by autoradiography. The gel was exposed to X-ray film.

The size of the extension products synthesized by Gerd_ePriS varied and could be very long. However, the addition of Gerd_ePriL drastically shortened the elongation products (Fig. 6A). The DNA primer on the M13 template was extended by ∼15 nucleotides by Gerd_ePriS but barely extended by Gerd_ePriSL in the presence of rNTPs under our experimental conditions. Similarly, the products of the extension of the M13-templated DNA primer by Gerd_ePriS were too large to enter the gel, and those by Gerd_ePriSL were much shorter (∼60 nt) in the presence of dNTPs.

In agreement with its ability to synthesize and extend a templated primer, Gerd_ePriS was capable of DNA binding (Fig. 6B). The apparent KD of the protein for the primed template used in the primer extension assay was 0.1∼0.25 mM, as estimated by the amount of the protein required for retarding half of the input DNA. In comparison, PriS (P41) from *Pyrococcus furiosus* is also able to bind DNA (Liu, et al. 2001) but that from *Saccharolobus islandicus* is not, consistent with the former being more closely related than the latter to the Heimdall group in the PriS evolutionary trees.

## Discussion

Phylogenetic analysis based on either PriS or PriL divides Asgard primases into two distinct groups, i.e., the Heimdall group and the Loki group. The Heimdall group enzymes cluster with eukaryotic primases, with the most closely related non-Asgard primases being those from thermophilic euryarchaea. The Loki group, represented by primases from Loki- and Thorarchaeota, with the most closely related non-Asgard primases being those from TACK superphylum. It is noticed that some branches of the primase-based phylogenetic trees do not agree with species classification. We speculate that the archaeal lineages represented by these branches may share similar habitats, or have present or past symbiotic or predatory relationships such that gene exchange has occurred at far greater efficiencies than the average level. This would have increased the diversity of archaeal primase, and ultimately driven the transition of archaeal primase toward eukaryotic primase. Additionally, no DNA polymerase α or subunit B, components of eukaryotic primase, were identified in the existing Asgard genomes.

The phylogenetic separation of Asgard primases is clearly supported by their structural differences. Notably, PriL of the Heimdall group, including Gerd_ePriL, resembles human PriL in containing additional three α-helices at the C-terminus. This unique C-terminal domain is not present in archaeal PriL outside the Heimdall group. This distinctive structural feature presumably allows enhanced protection of the conserved iron-sulfur cluster, increasing the stability of the enzyme, and, therefore, may have arisen as an evolutionary adaptation to the appearance of oxygen on the Earth.

Our biochemical assays revealed several unique properties of Gerd_ePriSL, a Heimdall group protein. Most significantly, Gerd_ePriSL synthesized very short primers (< 9 nt), which are much shorter than those produced by other well-characterized archaeal primases (e.g., *P. furiosus* and *S. solfataricus* P2 primases) (Liu, et al. 2001; Lao-Sirieix and Bell 2004; Lao-Sirieix, Nookala, et al. 2005). Short unit-length primers are typically produced by eukaryotic primases (Frick and Richardson 2001). It appears that primer size restriction might have been evolved in the Heimdall lineage of Asgard archaea. Gerd_ePriSL preferred dNTPs over rNTPS but required the presence of both types of nucleotides, with the optimal dNTPs:rNTPs ratio of 1:10, in primer synthesis in vitro. Further, although Gerd_ePriS alone showed detectable primer synthesis, this activity was enhanced, albeit slightly, by Gerd_ePriL. The ability to synthesize primers by PriS alone and the preferential use of dNTPs over rNTPs as the substrates in primer synthesis have been reported for other archaea, such as *P. furiosus* and *P. horikoshii* (Bocquier, et al. 2001; Matsui, et al. 2003).

Gerd_ePriSL, and Gerd_ePriS alone in particular, is also a robust DNA polymerase. Primer extension occurred with DNA primers and dNTPs, thus showing the same substrate preference as in primer synthesis. No significant primer extension was observed with RNA primers or rNTPs. Gerd_ePriL inhibited primer extension by Gerd_ePriS, reducing the size of the extension products to as short as ∼15 nt. By comparison, PriS from the phylogenetically close-related *P. furiosus* (*Pfu*P41) is also capable of independent primer extension, showing substrate preference for dNTPs (Bocquier, et al. 2001; Liu, et al. 2001), but PriS from the phylogenetically more distantly-related *S. solfataricus* P2 is unable to synthesize or extend primers on its own (Lao-Sirieix and Bell 2004; Liu, et al. 2015). On the other hand, the catalytic subunit P49 (PriS) of mouse primase is able to extend primers alone, producing short extension products (a few nucleotides), and the amount, but not the length, of the products was only slightly increased by the non-catalytic subunit P58 (PriL) (Copeland and Wang 1993). Taken together, our data suggest that the B18_G1 primase resembles primases from other archaea in some and eukaryotic primases in other biochemical properties. The ability of Gerd_ePriL to regulate the size of the products of primer synthesis and extension by the catalytic subunit Gerd_PriS is consistent with the need to minimize low fidelity synthesis in DNA replication, and presumably represents steps of evolution leading to the rise of eukaryotic primase.

In this study, we identified a three-residue motif, denoted motif G/S, in PriS. Motif G/S, which is typically Y/F/S*S in most archaeal primases and D/E*G in eukaryotic primases, allows distinction between archaeal and eukaryotic primases. However, motif G/S in PriS of the Heimdall group is Y*G, with the third residue identical to and the first residue different from the corresponding residue in the eukaryotic motif, an observation suggestive of the evolutionary intermediate stage of the enzyme. Statistical analysis reveals a total of five frequent amino acid substitutions, i.e., Y, F, D, S, and H, at the first position of motif G/S among all archaeal and eukaryotic primases. Since the latter four residues could readily be derived from Y though a single-base mutation, we speculate that Y is the ancestral residue at this site. Mutational analysis shows that the residue at this position influenced both the thermal stability and primer synthesis activity of Gerd_ePriS. Mutation of Y into F, a residue found in the *S. solfataricus* protein, lowered the activity of the enzyme, possibly an adaptation to slow growth often characteristic of archaea. Mutation of Y into D, a residue at the first position in motif G/S in the eukaryotic enzymes, reduced the thermal stability of the protein by ∼10°C but drastically enhanced the primase activity. The optimal temperatures for both primer synthesis and extension by Gerd_ePriSL are in the range of 50 to 55°C, suggesting that this strain may inhabit moderately hot environments. It is tempting to speculate that the Y-to-D substitution represents adaptation to cooling and increased growth efficiency during the evolution of eukaryotic ancestors (Lu, et al. 2024).

## Acknowledgements

We thank Wei Zhang (Institute of Microbiology, Chinese Academy of Sciences) for the technical assistance.

## Data availability

The data are available in the article or the online supplementary material.

