## Supplementary materials for "Phylogenetic and functional characterization of Asgard primases"

**This file includes:**

**Supplementary Figure 1-10**

**Supplementary Table 1-6**

A

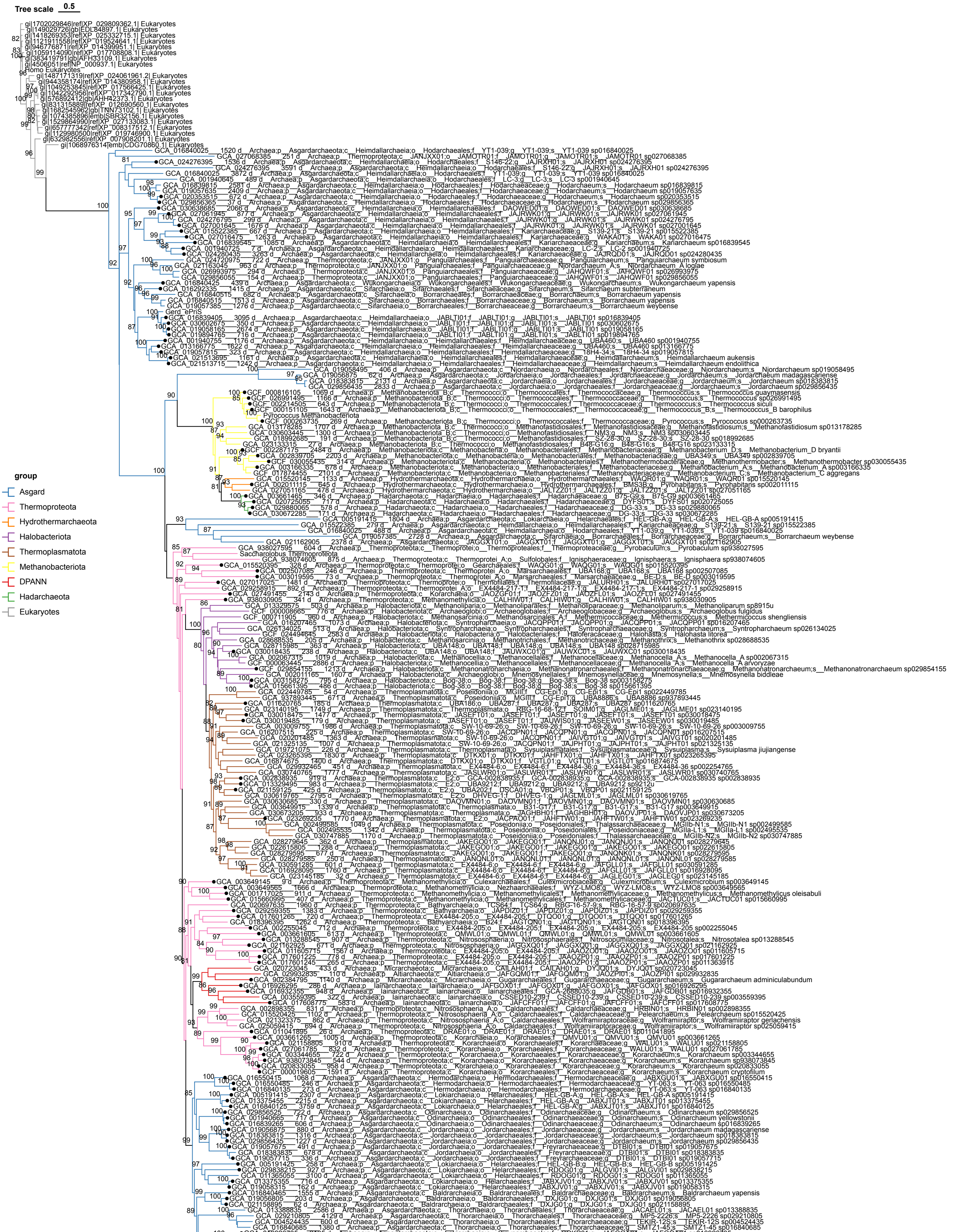
Tree scale 0.5
group
Asgard
Thermoproteota
Hydrothermarchaeota
Halobacteriota
Thermoplasmatota
Methanobacteriota
DPANN
Hadarchaeota
Eukaryotes

B

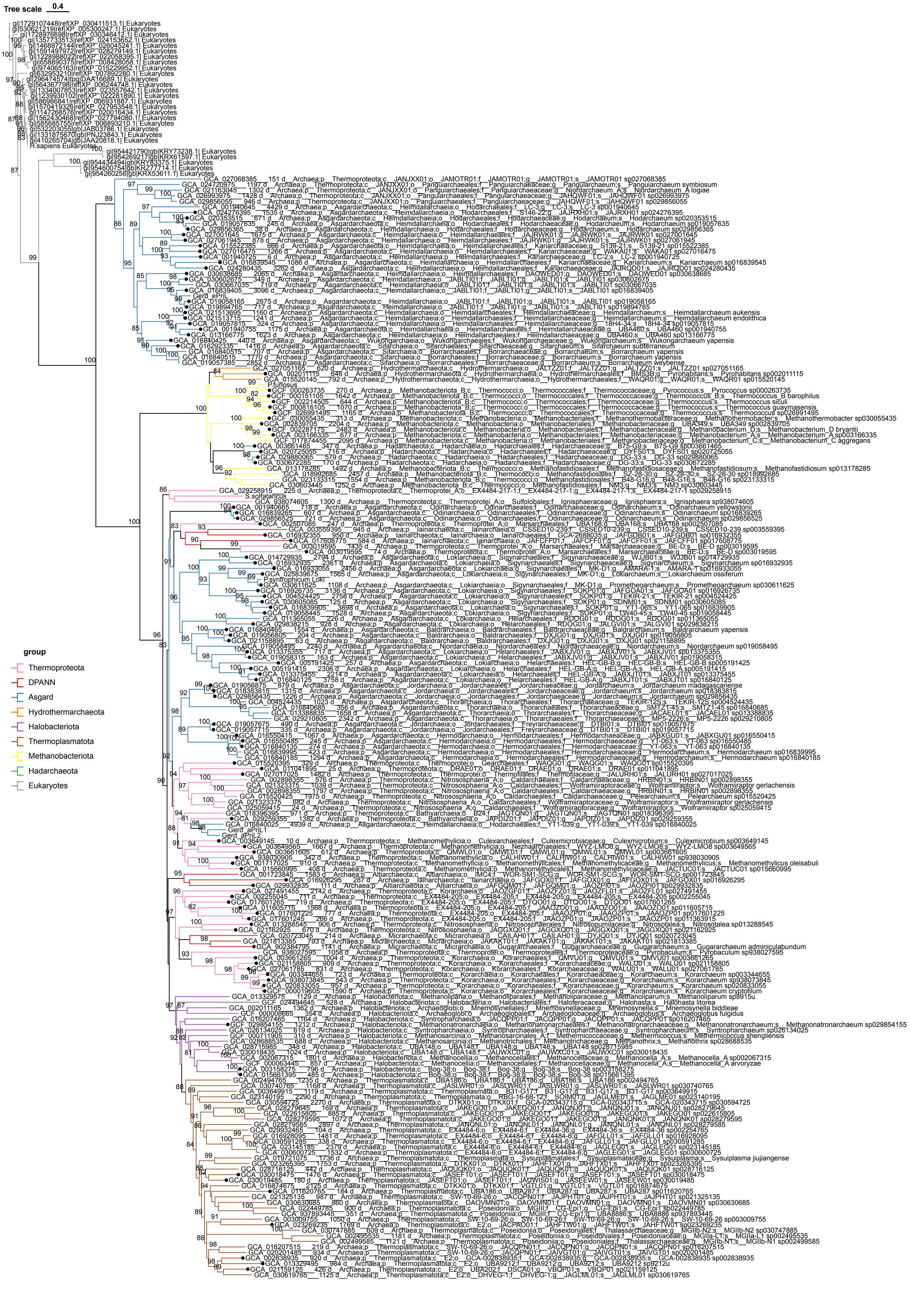

Tree scale 0.4
group
Thermoproteota
DPANN
Asgard
Hydrothermarchaeota
Halobacteriota
Thermoplasmatota
Methanobacteriota
Hadarchaeota
Eukaryotes

**Supplementary Figure 1. Phylogenetic trees of primases. (A)** Maximum likelihood phylogenetic tree of PriS. **(B)** Maximum likelihood phylogenetic tree of PriL. Detailed information of the proteins used to construct the phylogenetic tree is displayed in the figures. Solid dots indicate that genes encoding PriS and PriL are adjacently located on the genome.

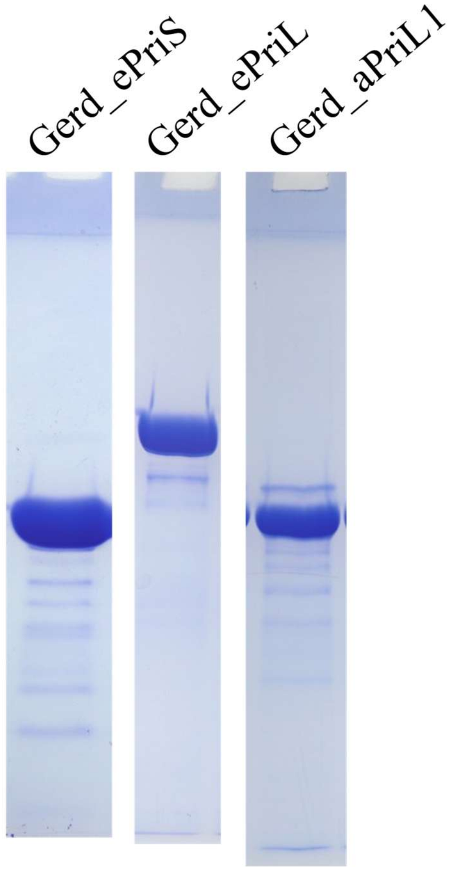


**Supplementary Figure 2. Purification of B18_G1 primase subunits.** Analysis of purified proteins by 12% SDS-PAGE.

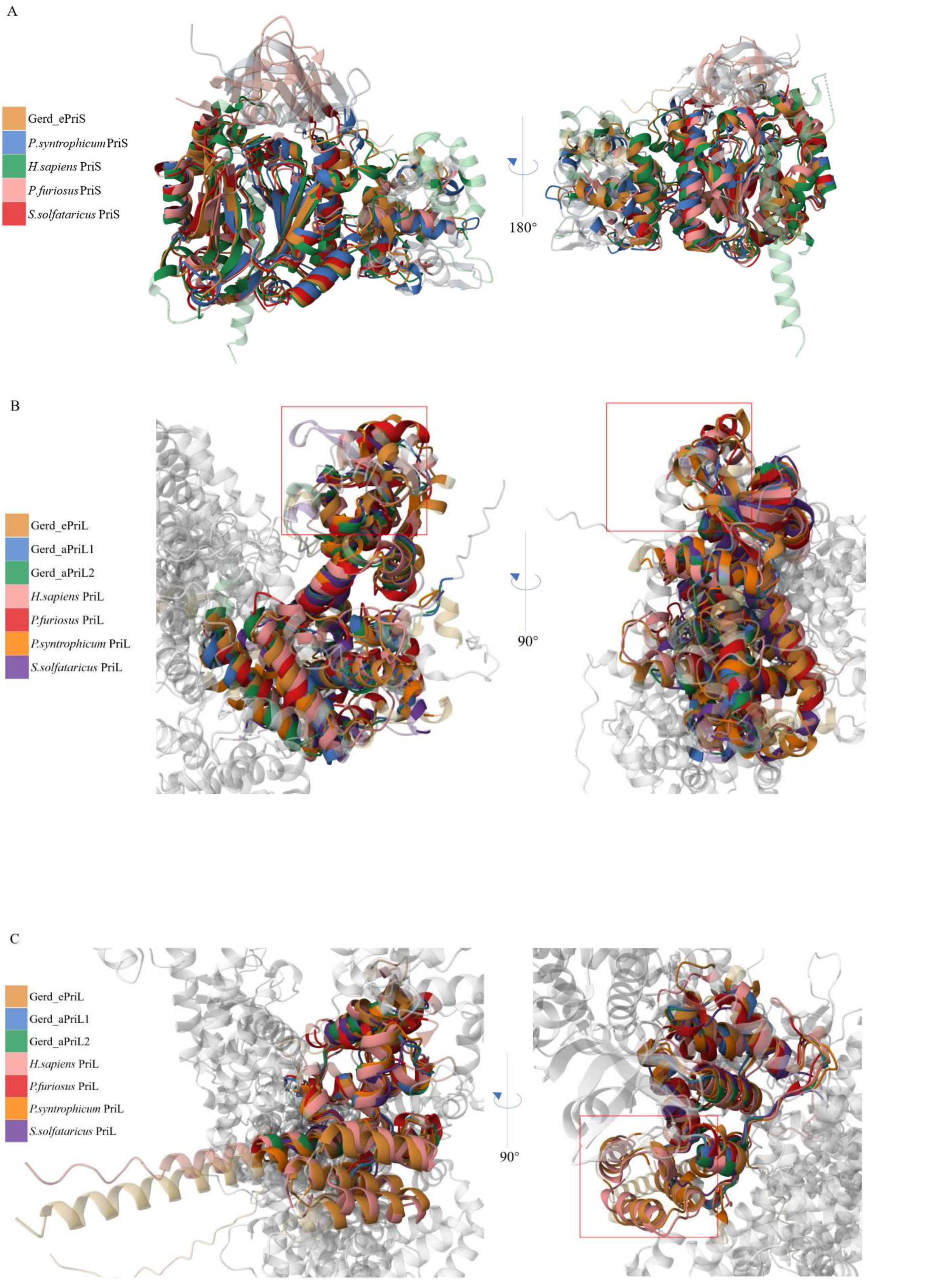


**Supplementary Figure 3. Structural comparison of selected archaeal and eukaryotic**

**primases. (A)** PriS. **(B)** The N-terminal domain of PriL. **(C)** The C-terminal domain of PriL. The structures of primases from *Candidatus* Gerdarchaeota archaeon B18_G1 (Gerd) and *Promethearchaeum syntrophicum* (*P. syntrophicum*) were predicted with AlphaFold, and those from human (*H. sapiens*; PDB: 4MHQ & 5EXR), *Saccharolobus solfataricus* (*S. solfataricus*; PDB: 5OF3), and *Pyrococcus furiosus* (*P. furiosus*; PDB: 1g71 & 9F28) were retrieved from RCSB PDB. The C-terminal part of PriL (Table S2) was predicted with AlphaFold3. Structurally distinct regions in PriLs of different origins are boxed. The colors corresponding to the different structures are indicated. The N-terminal and C-terminal domains of PriL were aligned in separate structural comparisons. The gray-white portion shows the C-terminal domain **(B)** or the N-terminal domain **(C)** of PriL. Structural alignments were performed using the Pairwise Structure Alignment tool on PCSB PDB with TM-align method.

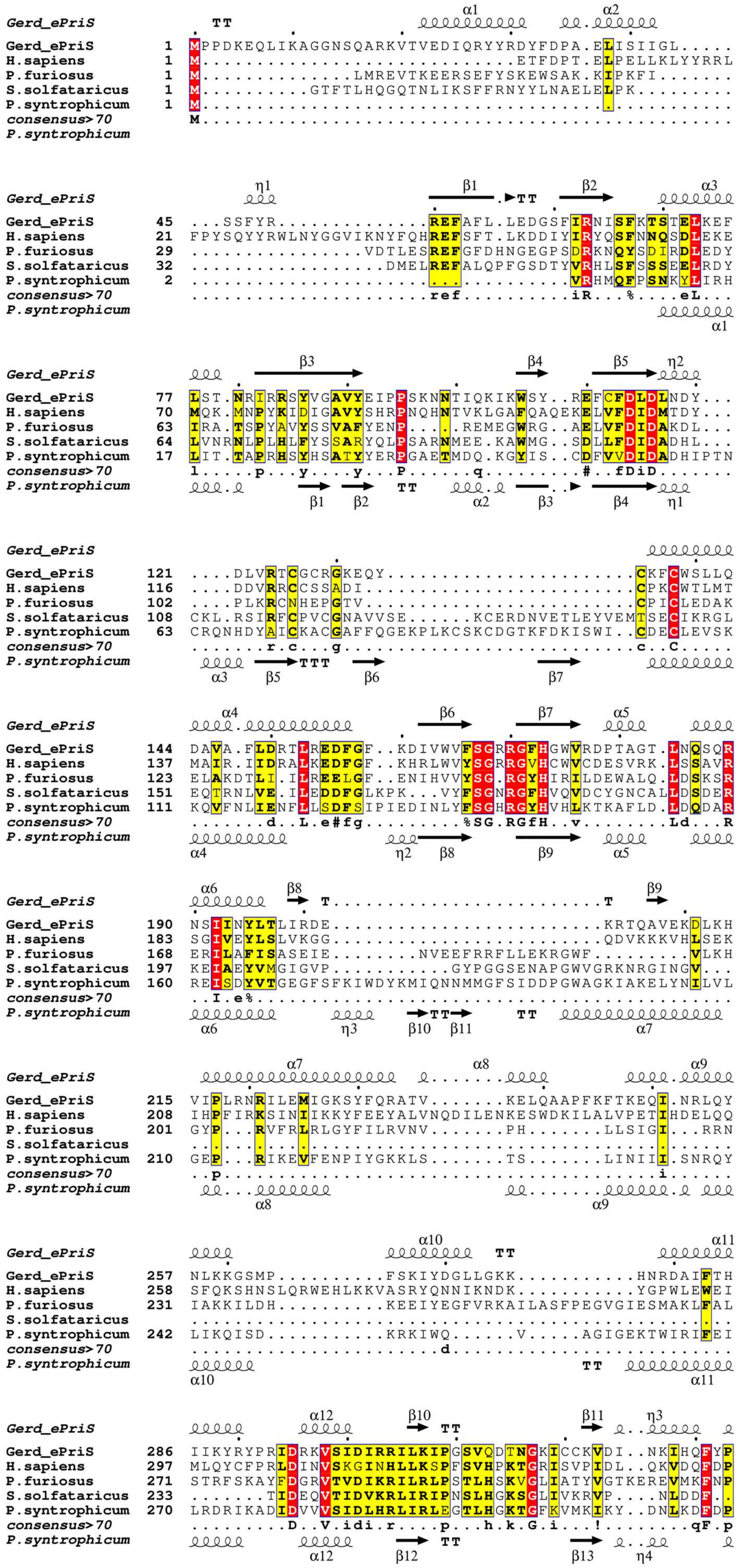

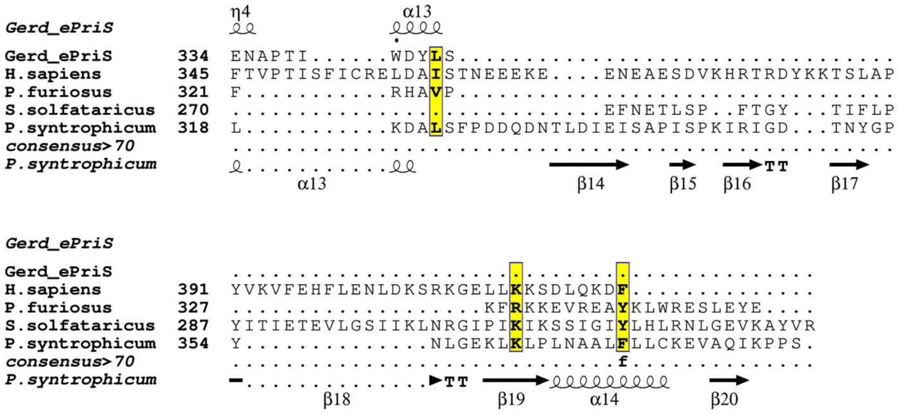


**Supplementary Figure 4. Secondary structure comparison of PriS.** The top panel shows the secondary structure diagram of Gerd_ePriS (squiggles for helices, arrows for β-strands and TT letters for turns), and the bottom panel displays that of PriS from *Candidatus Prometheoarchaeum syntrophicum* strain MK-D1. Secondary structure alignment was performed with ESPript 3.0. Identical and similar residues are boxed in red and yellow, respectively.

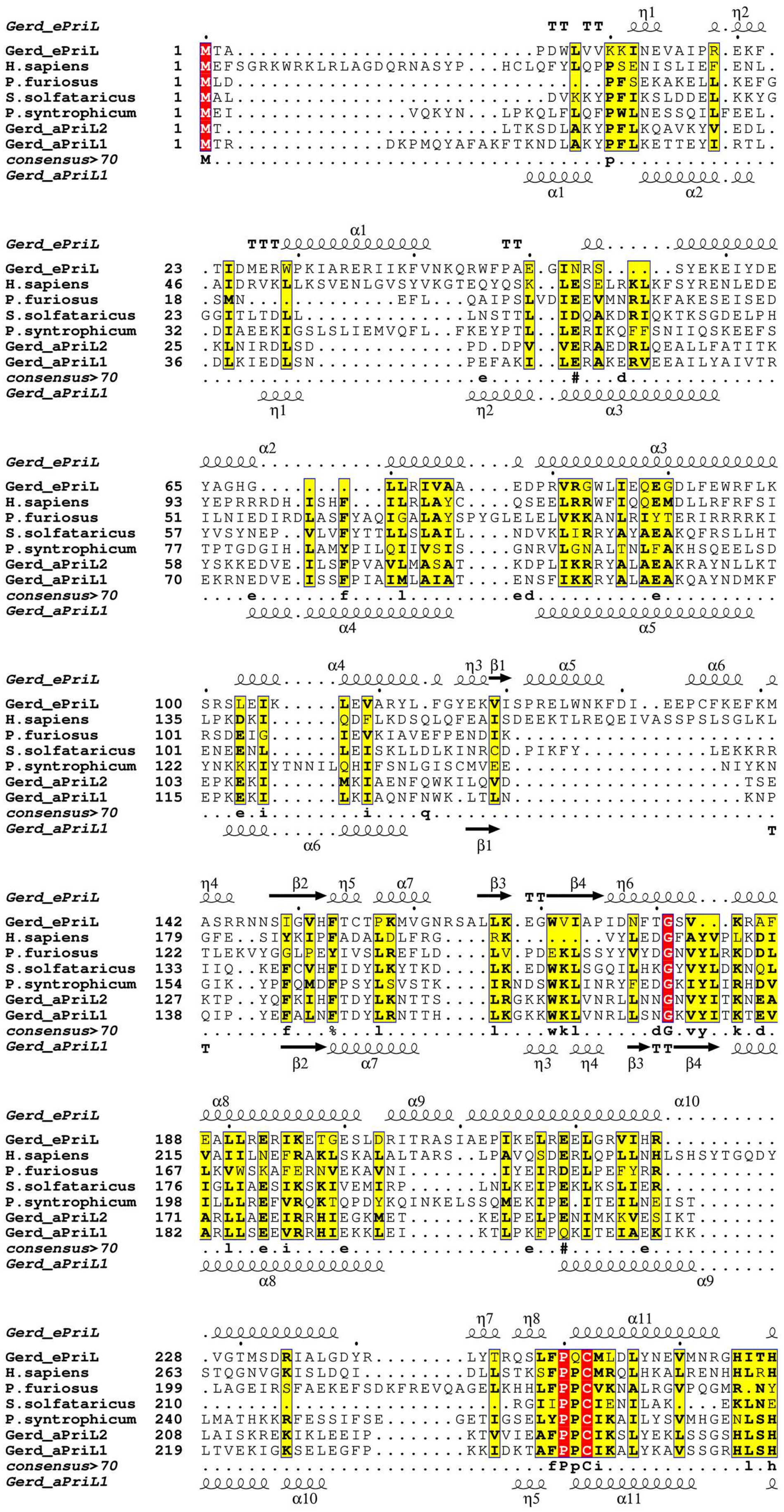
Gerd_ePriL
H.sapiens
P.furiosus
S.solfataricus
P.syntrophicum
Gerd_aPriL2
Gerd_aPriL1
consensus>70

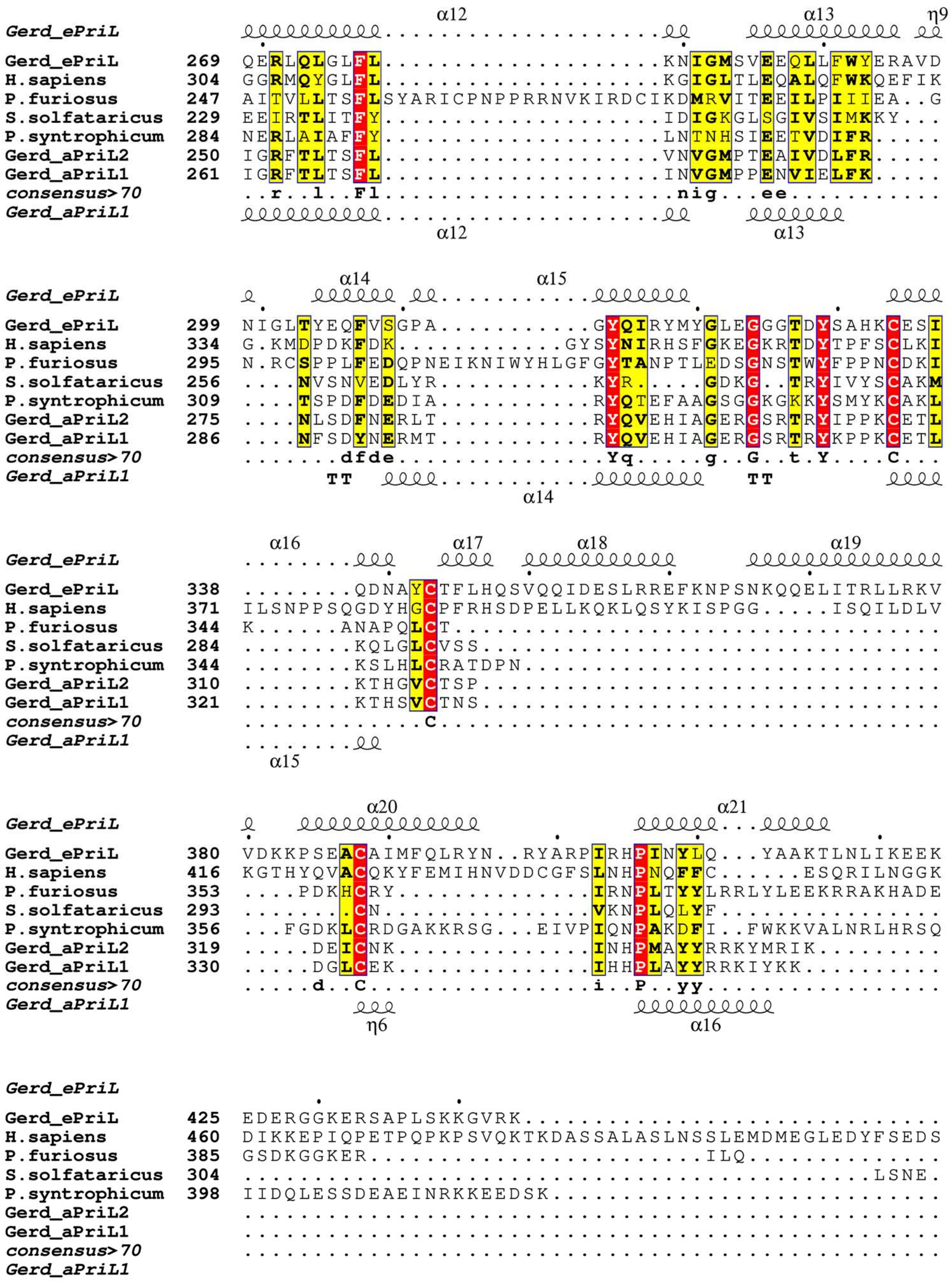


**Supplementary Figure 5. Secondary structure comparison of PriL.** The top panel shows the secondary structure diagram of Gerd_ePriL (squiggles for helices, arrows for β-strands and TT letters for turns), and the bottom panel displays that of Gerd_aPriL1. Secondary structure alignment was performed with ESPript 3.0. Identical and similar residues are boxed in red and yellow, respectively.

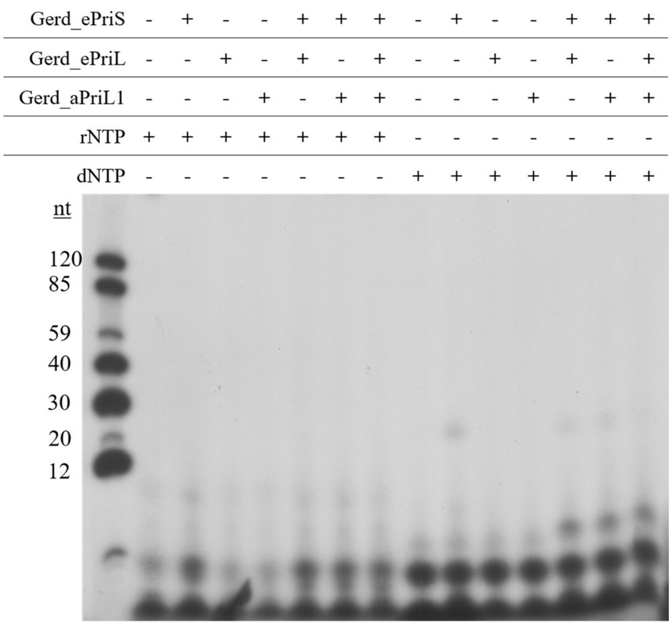


**Supplementary Figure 6. Primer synthesis by B18_G1 primase.** Reactions were performed by incubating single-subunit enzyme or multi-subunit complexes (1.5 μM) with DNA M13mp18 ssDNA (230 ng) and 10 μM dNTPs (1 μCi [α-$^{32}$P]dATP) or 10 μM rNTPs (1 μCi [α-$^{32}$P]dATP) in the standard assay mixture for 30 min at 55$^{o}$C. Reactions were stopped by the addition of SDS (0.8%) and protease K (1.6 mg/ml). The products were extracted with phenol/chloroform/isoamyl alcohol (25:24:1), precipitated with ethanol, and analyzed on 20% polyacrylamide gel (19:1) containing 8 M urea. The gel was exposed to X-ray film.

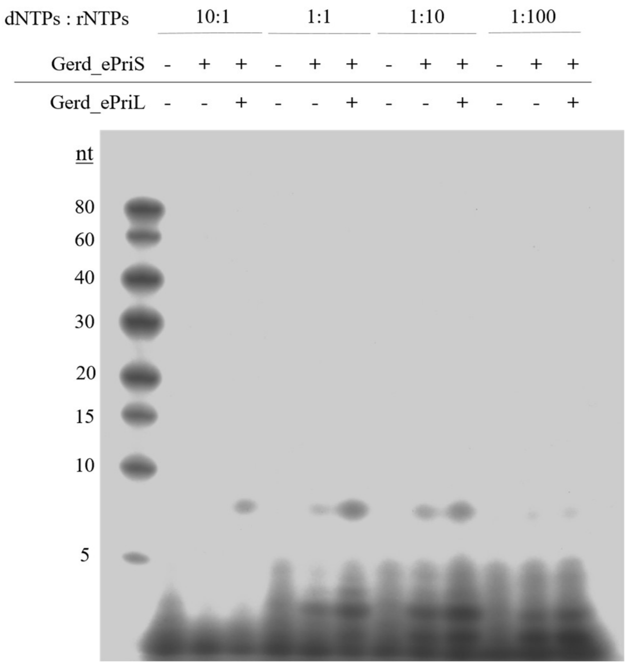


**Supplementary Figure 7. Primer synthesis by B18_G1 primase at various dNTPs/rNTPs ratios.** Reactions were performed by incubating Gerd_ePriS or Gerd_ePriS-ePriL (1.5 μM) with 230 ng M13mp18 ssDNA and different types of substrates in the standard assay mixture for 30 min at 55°C. The concentrations of dNTPs were 10 μM (1 μCi [α-32P]dATP). The concentrations of rNTPs were added according to the molar ratios shown at the top of the figure. Reactions were stopped by the addition of SDS (0.8%) and protease K (1.6 mg/ml). The products were extracted with phenol/chloroform/isoamyl alcohol (25:24:1), precipitated with ethanol, and analyzed on 25% polyacrylamide gel (19:1) containing 6 M urea. The gel was exposed to X-ray film.

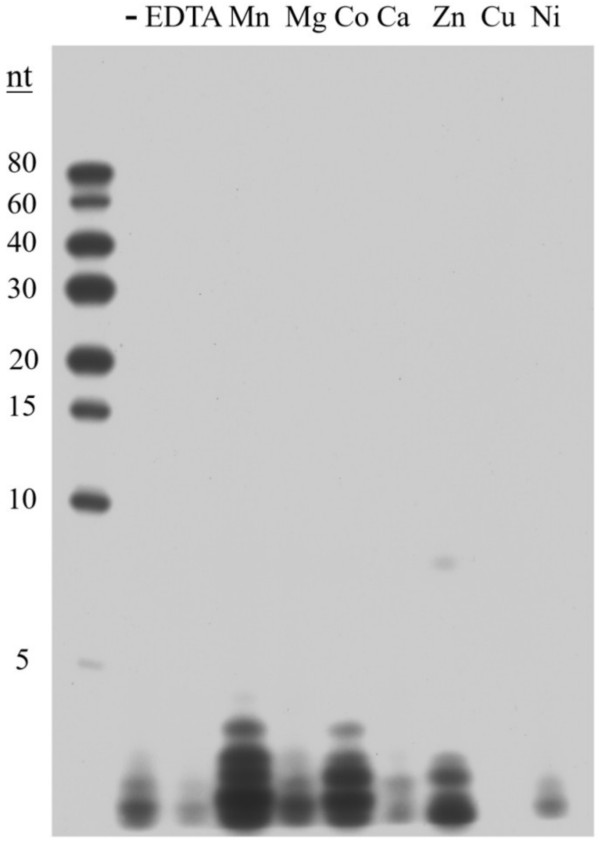


**Supplementary Figure 8. Effect of divalent cations on primer synthesis by Gerd_ePriSL.** The reaction mixture, containing Gerd_ePriS-ePriL (1.5 µM), M13mp18 ssDNA (230ng), 10 µM dNTPs (1 µCi [α-$^{32}$P]dATP) and 100 µM rNTPs, 50 mM MES-NaOH, pH7.0, 100 µg/ml BSA, and indicated divalent cations, was incubated at 55°C for 30 min. Reactions were stopped by the addition of SDS (0.8%) and protease K (1.6 mg/ml). The products were extracted with phenol/chloroform/isoamyl alcohol (25:24:1), precipitated with ethanol, and analyzed on 25% polyacrylamide gel (19:1) containing 6 M urea. The gel was exposed to X-ray film.

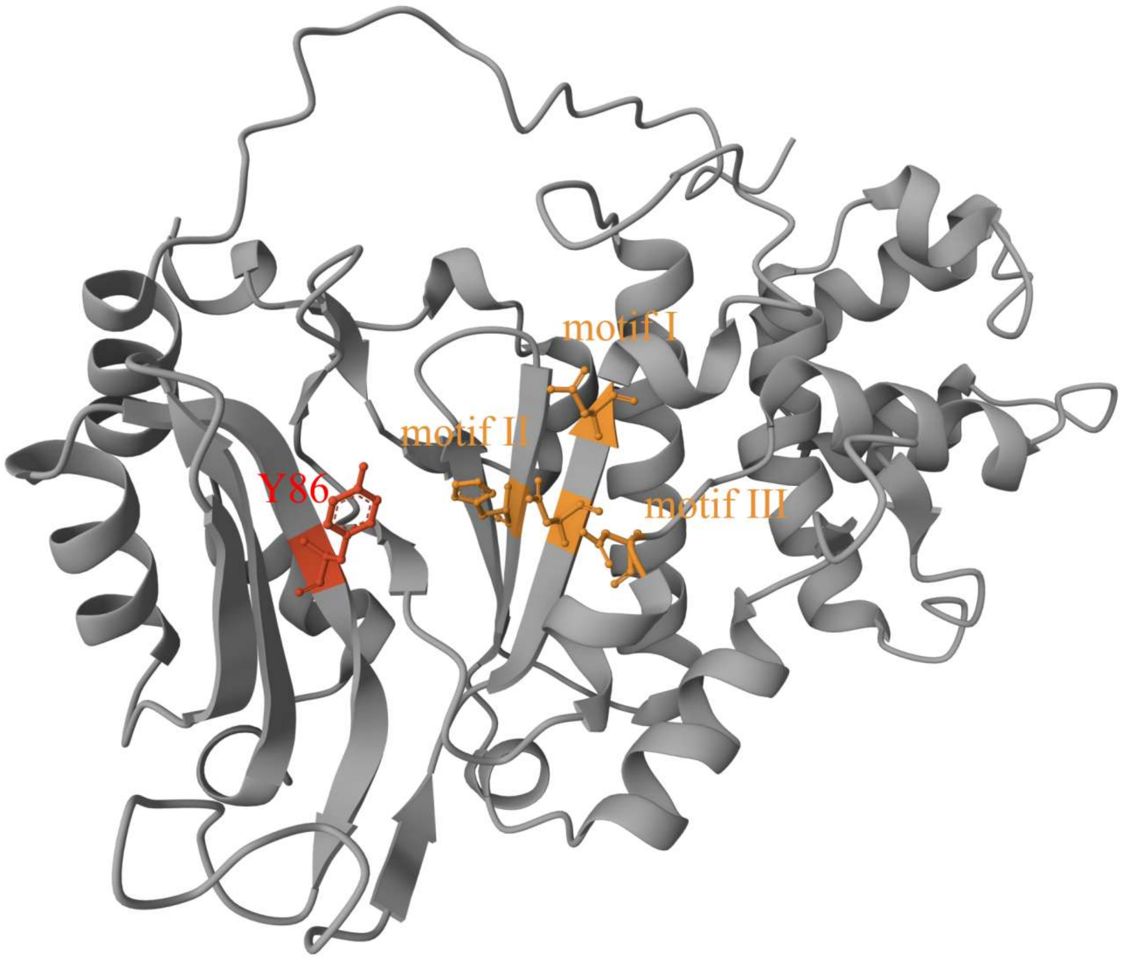


**Supplementary Figure 9. Schematic representation of the structure of Gerd_ePriS.** Motifs I, II, and III, the three conserved motifs in the AEP superfamily, are labeled with the conserved amino acid residues highlighted. Y86 is the first amino acid residue of motif G/S in Gerd_PriS, the site where point mutations were performed in this study. Structural visualization was performed by MolStar on RCSB PDB.

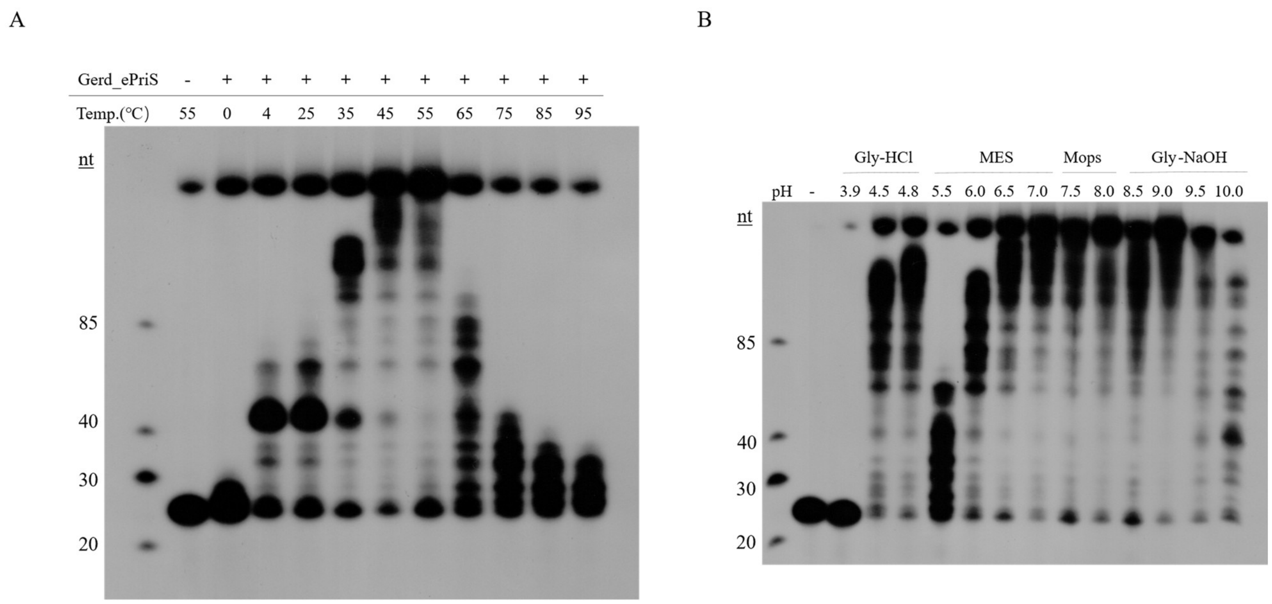


**Supplementary Figure 10. Effect of temperature and pH on primer extension by B18_G1 primase. (A)** Temperature. The reaction mixture, containing Gerd_ePriS (1.5 μM), 4 nM $^{32}$P-labeled D25 primer annealed to M13mp18 ssDNA, 10 μM dNTPs (1 μCi [α-$^{32}$P]dATP), 50 mM MES-NaOH, pH7.0, 100 μg/ml BSA, and 10 mM $MnCl_2$, was incubated for 30 min at indicated temperatures. **(B)** pH. Reactions were performed for 30 min at 55°C in the same mixture as in **(A)**, except for the substitution of Gerd_ePriSL for Gerd_ePriS, and in indicated buffers, instead of MES-NaOH. Reactions were stopped by the addition of SDS (0.8%) and protease K (1.6 mg/ml), and then analyzed on 15% polyacrylamide gel (19:1) containing 8 M urea. The gel was exposed to X-ray film.

**Supplementary table 1. Strains and oligonucleotides used in this study.**

| **species or fragment name** | **statement** | **sequences** |
|---|---|---|
| *Pyrococcus furiosus* | Genome DNA (Kept in our lab) | – |
| *Sulfolobus solfataricus* P2 | Strain and Genome DNA (Kept in our lab) | – |
| dT35 | ssDNA | 5’-TTTTTTTTTTTTTTTTTTTTTTTTTTTTTTTTTTT |
| SP2 | ssDNA | 5’TTTTTTTTTTTTTTTGTGTCGCAGCTGCCACCCTTTTT TTTTT |
| D25 | ssDNA | 5'-GTACCGAGCTCGAATTCGTAATCAT |
| R25 | ssRNA | 5'-GUACCGAGCUCGAAUUCGUAAUCAU |

**Supplementary table 2. The database IDs of the structures used in this study.**

| **Protein** | **ID** | **database** |
|---|---|---|
| PriS | | |
| Gerd_PriS | A0A497RBE9 | AlphaFoldDB |
| *Homo sapiens* PriS | 4MHQ | PDB |
| *Prometheoarchaeum syntrophicum* PriS | A0A5B9DFU6 | AlphaFoldDB |
| *Pyrococcus furiosus* PriS | 1g71 | PDB |
| *Saccharolobus solfataricus* PriS | 5of3 | PDB |
| PriL | | |
| Gerd_ePriL | A0A497R849 | AlphaFoldDB |
| Gerd_aPriL1 | A0A497QTN0 | AlphaFoldDB |
| Gerd_aPriL2 | A0A497QRQ5 | AlphaFoldDB |
| *Homo sapiens* PriS | P49643 | AlphaFoldDB |
| *Prometheoarchaeum syntrophicum* PriL | A0A5B9DG3 | AlphaFoldDB |
| *Pyrococcus furiosus* PriL | Q8U4H7 | AlphaFoldDB |
| *Saccharolobus solfataricus* PriL | Q9UWW1 | AlphaFoldDB |

**Supplementary table 3. Structure-resolved and -unresolved portions of PriL.**

| species (taxonomy) | PDB ID | resolved part | unresolved part |
|---|---|---|---|
| *Saccharolobus solfataricus* | 5OF3 | 3-266 aa & 293-306aa | 267-292 aa |
| *Pyrococcus abyssi* | 9F28 | 1-211 aa | 212-393 aa |
| *Pyrococcus horikoshii* | 2DLA | 1-222 aa | 213-394 aa |
| *Homo sapiens* | 5exr | 22-455 aa | 456-509 aa |

**Supplementary table 4. Information on structural alignment between Gerd_ePriS and PriS from other organisms.**

| Entry | RMSD | TM-score | Identity | Aligned Residues | Sequence Length | Modeled Residues |
|---|---|---|---|---|---|---|
| Gerd_PriS | - | - | - | - | 344 | 344 |
| *Promethearchaeum syntrophicum* PriS | 4.03 | 0.62 | 20% | 218 | 378 | 378 |
| *Homo sapiens* PriS | 3.09 | 0.81 | 30% | 284 | 425 | 400 |
| *Pyrococcus furiosus* PriS | 3.86 | 0.74 | 20% | 255 | 347 | 344 |
| *Saccharolobus solfataricus* PriS | 2.66 | 0.62 | 25% | 222 | 330 | 319 |

RMSD: Root Mean Square Deviation

**Supplementary table 5. Information on structural alignment between N-terminal portions of Gerd_ePriL and PriL from other organisms.**

| Entry | RMSD | TM-score | Identity | Aligned Residues | Sequence Length | Modeled Residue |
|---|---|---|---|---|---|---|
| Gerd_ePriL | - | - | - | - | 444 | 240 |
| Gerd_aPriL1 | 3.11 | 0.66 | 12% | 172 | 350 | 230 |
| Gerd_aPriL2 | 3.21 | 0.67 | 13% | 174 | 340 | 220 |
| *Homo sapiens* PriL | 3.52 | 0.7 | 14% | 181 | 509 | 260 |
| *Pyrococcus furiosus* PriL | 3.4 | 0.67 | 12% | 179 | 396 | 220 |
| *Promethearchaeum syntrophicum* PriL | 3.64 | 0.66 | 7% | 166 | 419 | 250 |
| *Saccharolobus solfataricus* PriL | 3.44 | 0.62 | 15% | 166 | 307 | 210 |

**Supplementary table 6. Information on structural alignment information between C-terminal domains of Gerd_ePriL and PriL from other organisms.**

| **Entry** | **RMSD** | **TM-score** | **Identity** | **Aligned Residues** | **Sequence Length** | **Modeled Residue** |
|---|---|---|---|---|---|---|
| Gerd_ePriL | - | - | - | - | 444 | 204 |
| Gerd_aPriL1 | 2.25 | 0.51 | 27% | 113 | 350 | 120 |
| Gerd_aPriL2 | 2.14 | 0.51 | 26% | 114 | 340 | 120 |
| *Homo sapiens* PriL | 2.37 | 0.77 | 27% | 164 | 509 | 250 |
| *Pyrococcus furiosus* PriL | 3.29 | 0.48 | 21% | 107 | 396 | 176 |
| *Promethearchaeum syntrophicum* PriL | 3.24 | 0.54 | 24% | 117 | 419 | 169 |
| *Saccharolobus solfataricus* PriL | 2.11 | 0.42 | 17% | 94 | 307 | 97 |
